# Bighorn sheep T2T genome assembly reveals differences in immune genes: a potential cause of high morbidity due to respiratory pathogens

**DOI:** 10.1101/2025.09.30.679298

**Authors:** Temitayo A. Olagunju, Mariia Pospelova, John C. Schwartz, Michelle R. Mousel, Lindsay M.W. Piel, Paige C. Grossman, Kathryn P. Huyvaert, Kristen L. Kuhn, Tajbir Raihan, Morgan R. Stegemiller, Sarem F. Khilji, Gordon K. Murdoch, Ahmed Tibary, Lisette P. Waits, Arang Rhie, Sergey Koren, Adam M. Phillipy, Stephanie D. McKay, Shannon M. Clarke, Emily L. Clark, Rudiger Brauning, Noelle E. Cockett, John A. Hammond, Maggie Highland, Yana Safonova, Timothy P.L. Smith, Benjamin D. Rosen, Brenda M. Murdoch

## Abstract

The bighorn sheep (*Ovis canadensis*), despite its close relation to domestic sheep, suffer higher morbidity and mortality from respiratory disease complexes, likely due to genetic differences in immune responses. Unraveling highly repetitive regions such as immune loci and genetic differences was problematic until now. We generated a bighorn sheep telomere-to-telomere assembly, adding 14.28% of novel sequence compared to the previous reference. This enabled the first complete immune loci annotation revealing the IGL and TR loci are significantly short in bighorn sheep. Importantly, a critical immune gene *GBP5* and *ZNF501*, involved in Golgi-mediated immune response, are lacking in bighorn but present in domestic sheep. Re-analysis of a *Mycoplasma ovipneumoniae* carriage study, using this assembly, identified the immune gene *CAPN2* as a key genetic marker for disease carriage, not observable in the original study. This work provides a critical resource for identifying phenotype-linked genetic variation and exploring evolutionary adaptations of bighorn sheep.

## INTRODUCTION

North American bighorn sheep (*Ovis canadensis,* OCA), diverged from domestic sheep (*Ovis aries*, OAR) approximately 5.3 million years ago^1^, and are an iconic species of deep cultural and economic importance for Native Americans^2^. Once numerous, their populations have been decimated due to habitat loss, overhunting, and, importantly disease, an issue that continues to affect bighorn sheep populations today ^3–5^. In particular, a key pathogen that contributes to respiratory disease in bighorn sheep, *M. ovipneumoniae*, may be transmitted from chronic carriers to naïve individuals within bighorn sheep herds, domestic sheep, goats and other wildlife^5,6^. While domestic sheep are also known to suffer from pneumonia, the severity and mortality are notably higher in bighorn sheep^7^. The cause of these disparate outcomes remains a subject of continuous scrutiny, and the underlying genetic factors are yet to be fully elucidated^7,8^.

Investigating the genetic basis for this differential susceptibility has been hampered by the lack of a high-quality reference genome. Previous studies have relied on a highly fragmented bighorn genome assembly (GCA_004026945.1) that could not resolve the complex architecture of critical immune loci^7,8^. These loci are notoriously difficult to assemble due to their polymorphic and repetitive nature.

The innate immune system provides an immediate generalized response to cellular damage, microbial components and certain non-self proteins, the adaptive immune system develops specialized responses by producing T-cell receptors (TRs) and antibodies that can bind specific antigens^9,10^. The innate immune system uses germline-encoded pattern recognition receptors to detect molecular patterns which are damage-and pathogen-associated. Damage or introduction of a pathogen triggers rapid inflammatory responses through innate immune cells such as macrophages, neutrophils, dendritic cells, and natural killer (NK) cells^9,10^. The genomic complexity at the adaptive immune loci is enhanced by two main factors; (i) TRs and antibodies are not directly encoded in the genome but through somatic V(D)J (variable, diversity and joining) gene recombination of germline TRs and immunoglobulin (IG) loci^11,12^. (ii) The major histocompatibility complex (MHC) bridges the innate and adaptive immunity through the ability to distinguish self and non-self antigens^13^. It is classified into class I (MHC-I) and class II (MHC-II) genes, and the MHC-I region in bovids and other artiodactyls is split across two regions^14^. Previous studies of mammalian adaptive immune loci have also revealed that, on average, IG loci are more diverged compared to TR loci^15,16^, possibly due to evolutionary constraints related to binding to MHC. Successful assembly and characterization of these polymorphic and repetitive but highly critical genomic regions thus require contiguous T2T whole genome assemblies. Recent success of telomere-to-telomere (T2T) whole genome assembly in mammals^15,17,18^ provides a powerful opportunity to finally overcome these limitations and enable high-resolution exploration of immune diversity in bighorn sheep.

In this study, we present a complete, chromosome-scale T2T genome assembly of bighorn sheep (GCF_042477335.2, *ARS-UI_OviCan_v2* on NCBI; Bighorn-T2T, henceforth). Our assembly facilitated the first comprehensive annotation of both innate and adaptive immune loci in this species and a direct comparison to the domestic sheep. To demonstrate its utility, we show that Bighorn-T2T provides a superior framework for the identification of genetic variants associated with *Mycoplasma ovipneumoniae* disease carriage compared to using a domestic sheep reference. This resource is an invaluable tool for deepening our understanding of the differential outcomes of infection by respiratory disease pathogens in bighorn and domestic sheep.

## RESULTS

### Genome sequencing and T2T assembly

A mix of short reads and long reads (HiFi and nanopore) sequence data were generated from tissues of a male and a female fetus from Rocky Mountain Bighorn sheep sires and Polypay domestic sheep dams inter-species crosses for assembly (Extended data Fig.1). The Verkko assembler^19^ v2.0 produced 19 single-contig T2T chromosomes of the paternal bighorn haplotype (from the male fetus) while the remaining eight chromosomes were subjected to additional manual curation steps to bring them to telomere-to-telomere status (Supplementary Data 1, Supplementary Methods). The X chromosome was assembled T2T in one contig from the female fetus and was added to the male haploid assembly to produce the complete bighorn sheep T2T assembly (Bighorn-T2T) containing all the autosomes and both sex chromosomes (Figure 1a, Table 1, Extended data Fig.1). Assembly completeness evaluation with Compleasm^20^ recovered 98.31% complete BUSCOs (basic universal single-copy orthologous genes) from the *cetartiodactyla_odb10* database (Supplementary Data 2) while 63.76 QV base accuracy was estimated by Merqury^21^ (Supplementary Data 3) after manual curation and polishing.

**Figure 1.**
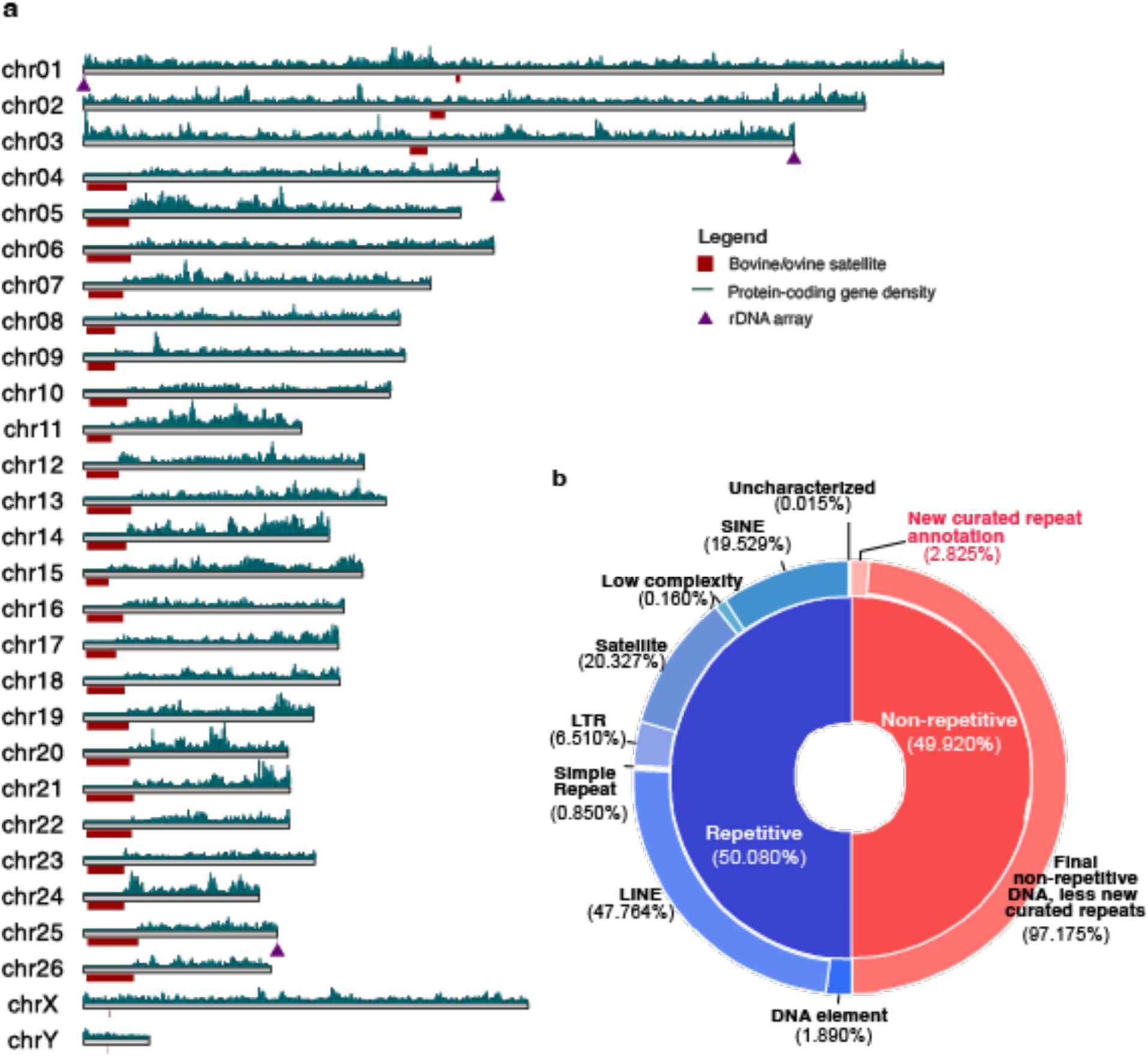
Structure of the complete Bighorn-T2T genome assembly highlighting the genomic features and repeat content. (a) The complete bighorn sheep assembly showing the bovine/ovine satellites enriched at the (peri)centromeric region (red blocks below the chromosomes). The rDNA array loci are highlighted as purple triangles below the chromosomes. The protein-coding genes densities are shown above the chromosomes highlighting the acrocentric p-arms lacking protein-coding genes. (b) The repetitive (blue) and non-repetitive (red) DNA content of the bighorn sheep genome showing the proportion of the repeat content before manual repeats curation in the inner sector (repetitive is blue, 50.080% and non-repetitive is red, 49.920%). The distribution of the content of each inner sector is displayed in the outer segments. The outer segment of the non-repetitive (red) half shows the additional portion that was finally annotated as repetitive with manual repeats curation.

**Table 1:**
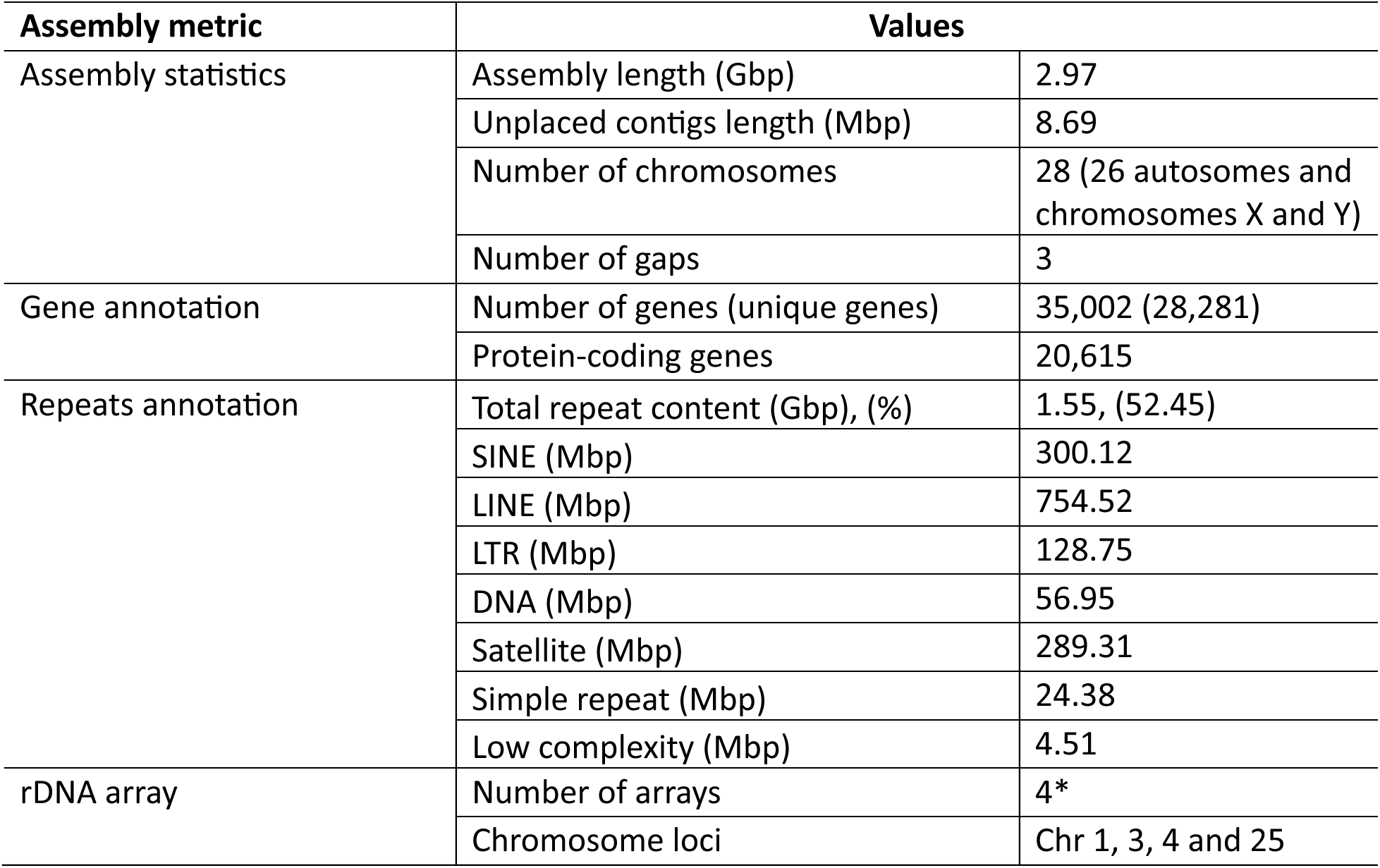
Bighorn-T2T genome assembly metrics.

SINE, short interspersed nuclear element; LINE, long interspersed nuclear element; LTR, long terminal repeat. *Five arrays were expected according to a previous report on domestic sheep nucleolar organizer region (NOR) based on silver staining (see details in supplementary methods)

### Annotation of genomic features

More than half of Bighorn-T2T (52.90%) was repetitive DNA (Figure 1b, Supplementary Data 4), comprised predominantly of LINE elements as observed in human and mouse^22,23^. The bovine satellites characterizing the centromeres, identifiable as a highly homogenized sequence signature on a chromosome self dot plot (Figure 2a), were found to be longer (7.2Mb - 16.2Mb) on the acrocentric autosomes (chromosomes 4-26) than the 1.1Mb-5.7Mb range on the submetacentric autosomes (chromosomes 1-3) (Figure 2b, Extended Data Fig.2, Supplementary Data 1). A previously unidentified higher-order repeat (HOR) sequence spanning 31.12Mb of the genome was newly annotated and observed to be enriched at most of the centromeres (Figure 2a, Extended Data Fig.2, Supplementary Data 5 and 6). Segmental duplications (seg dups, defined as genome segments with >= 1kb length and >= 90% sequence identity) comprising 9.49 Mb and 7.08 Mb intrachromosomal and inter-chromosomal seg dups, respectively, covered 16.57Mb (0.56%) of the bighorn sheep genome (Supplementary Data 7).

**Figure 2.**
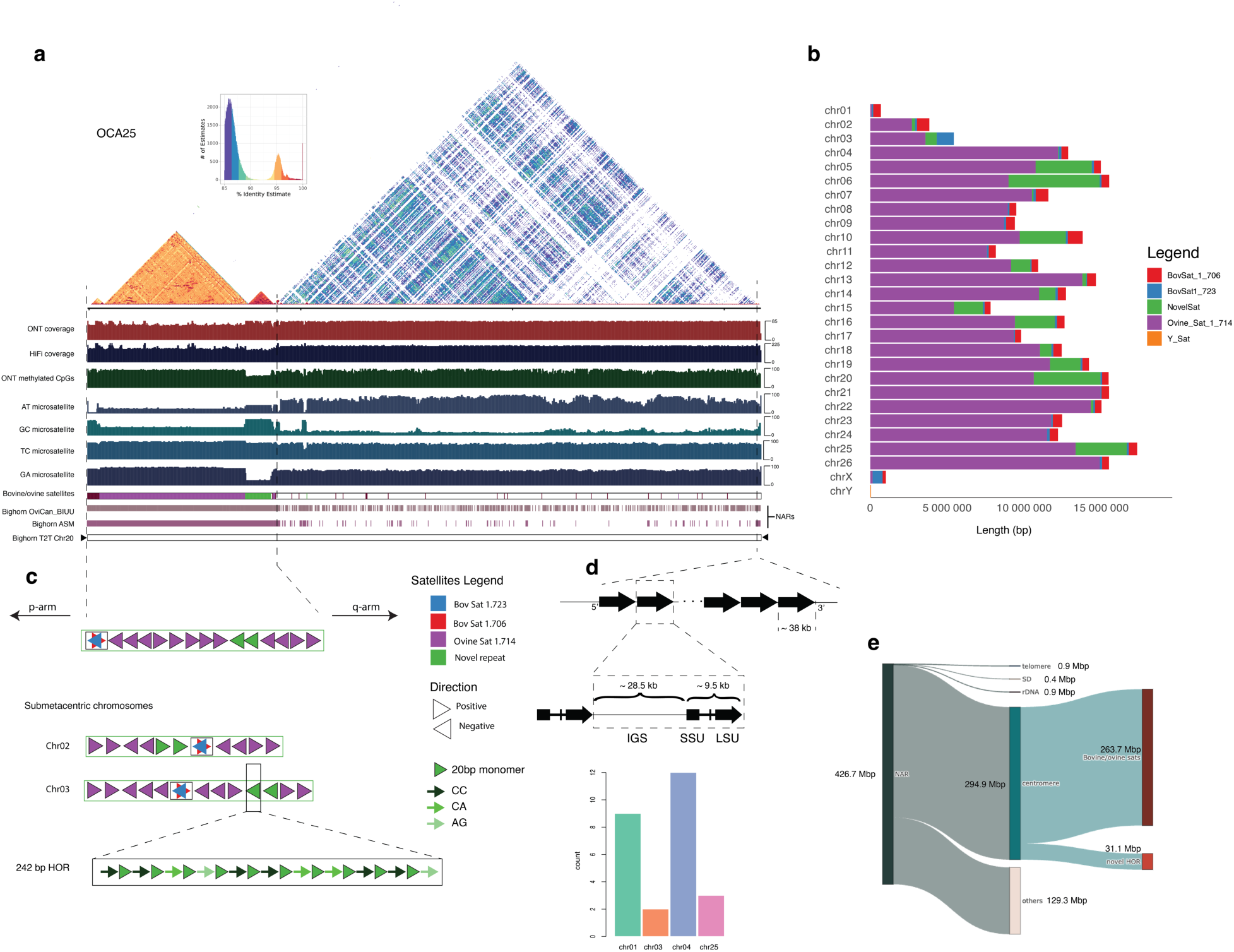
Overview of a Bighorn-T2T chromosome (OCA25) (a) The self dot plot of a representative acrocentric autosome (OCA25) highlighting the highly repetitive sequence content from the centromeric region up to the end of the short p arm where the yellow/deep red triangular segment corresponds to homogenized satellite sequence classes; ONT reads coverage track; HiFi reads coverage track; ONT methylated CpGs; AT, GC, TC and GA microsatellite tracks are data quality check against known sequencing technology biases in repetitive regions. The Bovine/ovine satellites track shows the loci of the centromere-enriched sequence as well as a previously unidentified higher-order repeat enriched at the centromere. Other tracks show the newly assembled regions (NARs) relative to the two previously available bighorn sheep genome assemblies on NCBI and the Bighorn-T2T with the telomeres represented with the black triangles. (b) The distribution of and coverage of the different satellite sequence classes enriched at the centromere. (c) The content and organization of the centromeric region of the acrocentric chromosome (followed by the submetacentric chromosomes 2 and 3) highlighting the enrichment of the bovine satellites 1.723 and 1.706 which are co-located but are interlaced with one another on opposite strands, the ovine satellite 1.714 which is reported to characterize and are enriched at the centromeres of ovine species, and a newly identified higher order repeat (HOR) sequence. The HOR is comprised of a 20bp monomer which is organized as tandem copies of the monomer interlaced with the dinucleotides CC, CA and AG as shown. (d) The structure of the rDNA unit organized as a tandem array on the long arm of the acrocentric chromosome comprising the intergenic spacer (IGS), short sub-unit (SSU) and the long sub-unit (LSU) annotated by Repeatmasker using the human homolog. The number of copies annotated on each chromosome is shown in the barplot. (e) Distribution of the genomic features and sequence classes of the newly assembled regions (NARs) of Bighorn-T2T relative to the older bighorn sheep assemblies GCA_004026945.1 and GCA_001039535.1 on NCBI.

NCBI RefSeq structural annotation of Bighorn-T2T identified 35,002 (20,615 protein-coding and 14,387 non-coding) genes in addition to 4,512 pseudogenes. The short p-arms of the acrocentric chromosomes lacked protein-coding genes but harbored a few non-coding genes (Supplementary Data 1). On the sex chromosomes, the recombinant pseudo autosomal regions (PAR) of the bighorn sheep chromosome X (OCAX, 7,085,862 bp) and chromosome Y (OCAY, 7,067,061 bp) exhibited consistent organization and content with mammals^24,25^ except for five genes exclusive to both sex chromosomes (Supplementary Data 8). The rDNA array-bearing NORs were located on OCA1, OCA3, OCA4 and OCA25 of Bighorn-T2T, (Figure 1a, Figure 2d) contrary to the reported arrays on OAR1, OAR2, OAR3, OAR4, and OAR25 on the domestic sheep based on silver-staining of the domestic sheep genome^26^. This difference is due to the absence of an rDNA array on OCA2 of the male fetus (Supplementary methods).

### Newly assembled bighorn sheep genomic regions

The previous bighorn sheep reference assembly on NCBI (GCA_004026945.1, Bighorn-v1 henceforth) was a 2.9Gb scaffold-level assembly (comprising 1,048,136 scaffolds with scaffold N50 and L50 of 69.4kb and 11,415 respectively) which lacked any telomere sequence on any of the scaffolds (Supplementary Data 9). Bighorn-T2T added 14.28% (426.70Mb) of NARs (Figure 2e) in comparison to Bighorn-v1 (Supplementary Data 10). The NARs include 39.63% of OCAY, comprising mainly the highly repetitive ampliconic region (Supplementary Data 10). A total of 122 genes (34 protein-coding and 88 non-coding) were not observed on Bighorn-v1 (Supplementary Data 11) and newly added to Bighorn-T2T. Some protein-coding genes of interest in this set include *LOC138437835* (a C-type lectin domain family 2 member D11-like; a paralog of *CLEC2D*) located within the highly complex natural killer cell region on OCA3; and *LOC138419301* (putative killer cell immunoglobulin-like receptor like protein KIR3DP1) in the leucocyte receptor complex on OCA14.

### Bighorn and domestic sheep genome comparison

The OCAX was 2.20Mb longer than OARX but 193.5 kb shorter than the X chromosome of a domestic sheep T2T assembly^27^, OARX-T2T (Figure 3a) (Supplementary data 12). Inversions were observed between OCAX and OARX that were not observed in the alignment to OARX-T2T (Figure 3a). Similarly, OCAY was about 4Mb shorter than OARY^28^ (21.49 Mb vs. 25.91Mb) (Figure 3a) because of an expansion of ampliconic gene families on OARY (Supplementary Data 13).

**Figure 3.**
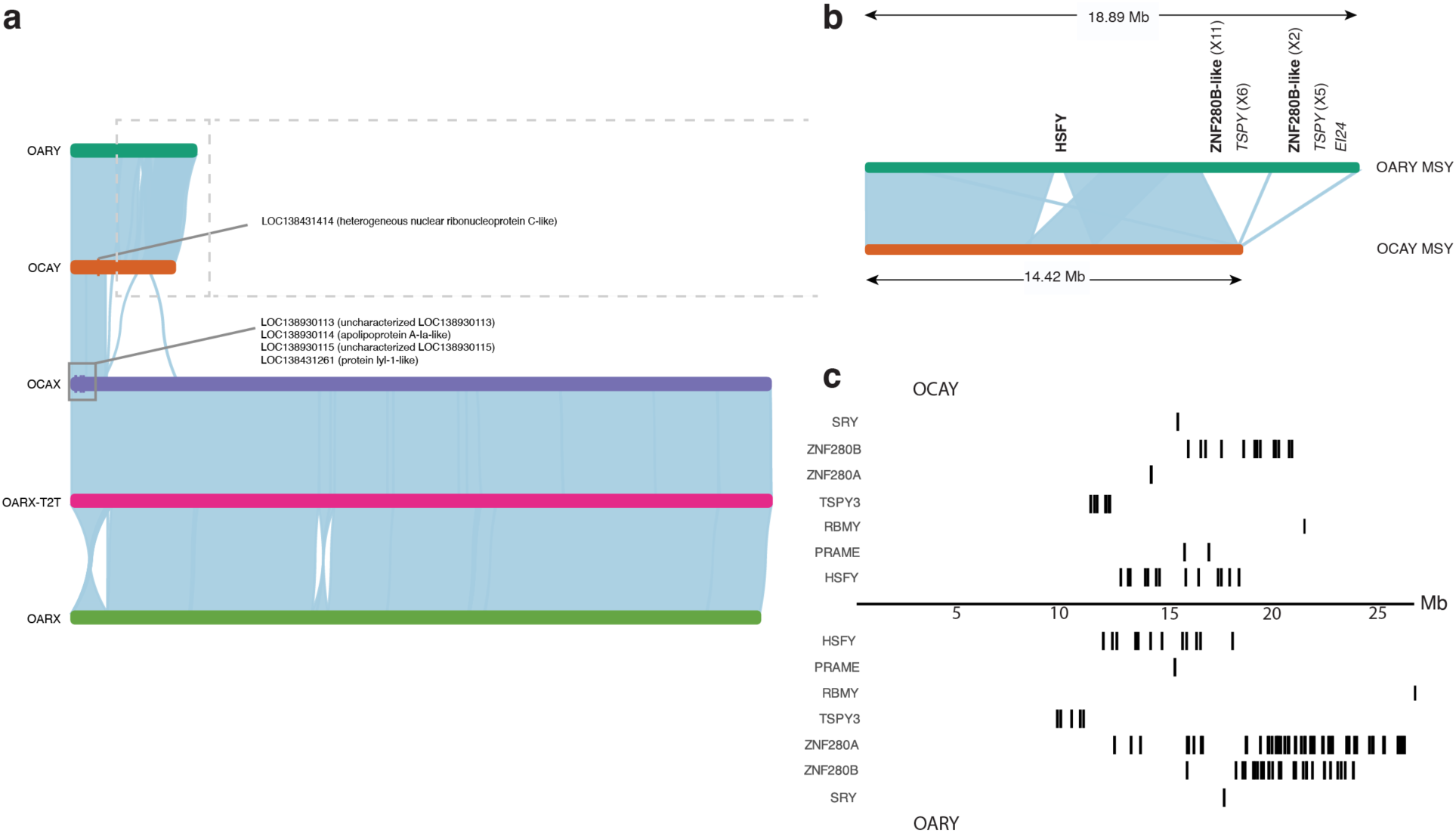
**Bighorn and domestic sheep comparison reveal structural differences in the sex chromosomes**. (a) Alignment of OARY, OCAY, OCA X, OARX-T2T, and OARX, respectively. General collinearity is observed between the sex chromosomes except two large-scale inversions between OARX and OARX-T2T and OCAX; A 7.39Mb inversion was observed between OCAX and OARX in the recombinant pseudo-autosomal region (PAR) with breakpoint around the centromeric satellite repeat of OCAX, and a 3.30 Mb inversion was observed at 50.02 Mb and at 51.55 Mb on OCAX. The PAR of OARX is inverted relative to OARX-T2T and OCAX. (b) Zoomed in view of the MSY of OARY and OCAY showing 4.4Mb extra DNA on OARY MSY region relative to OCAY. The protein-coding genes (bold font) and pseudogenes (light font) with the number of copies (in parentheses) in the region of extra DNA on OARY are shown. There were more ampliconic genes on OARY MSY than on OCAY MSY. (c) Loci of *SRY* gene and the ampliconic genes family on OCAY and OARY.

Three alignment gaps totaling 6.36Mb between the male-specific Y (MSY) region of OCAY and the longer OARY harbored 14 more protein-coding genes and 12 more pseudogenes on OARY (Figure 3b, Supplementary Data 13).

Structural differences between Bighorn-T2T and the domestic sheep genomes revealed 69.05Mb (Figure 4a, Supplementary Data 14) of predominantly large insertions and deletions (INDELs, length >= 50bp) (Figure 4b, Supplementary Data 14). SnpEff^29^ variant effect predictor predicted that a total of 9,828 genomic features were affected by these genomic differences (Supplementary Data 15). Out of these, 950 features were annotated as having high impact. The subset with the highest number of variants (≥3) in the high impact category contained ten protein-coding genes (and one pseudogene) with immune-related functions (Supplementary Data 15). These genes include *BIN2* (bridging integrator 2), *CTPS1* (CTP synthase 1), *ENDOU* (endonuclease, poly(U) specific), *PPM1A* (protein phosphatase, Mg2+/Mn2+ dependent 1A), *LOC101123578* (leukocyte immunoglobulin-like receptor subfamily A member 6), *LOC114108841* (immunoglobulin lambda-1 light chain), the pseudogene *LOC114114065* (killer cell lectin-like receptor subfamily B member 1), *CALR* (calreticulin), *IFI208-like* (Interferon-activable protein 208-like), *BTN2A1-like* (butyrophilin subfamily 2 member A1-like) and *MUC5B* (mucin 5B, oligomeric mucus/gel-forming). The significant effect of the structural differences on these genes was predicted due to exon loss and/or frameshift variant (Supplementary Data 15).

**Figure 4.**
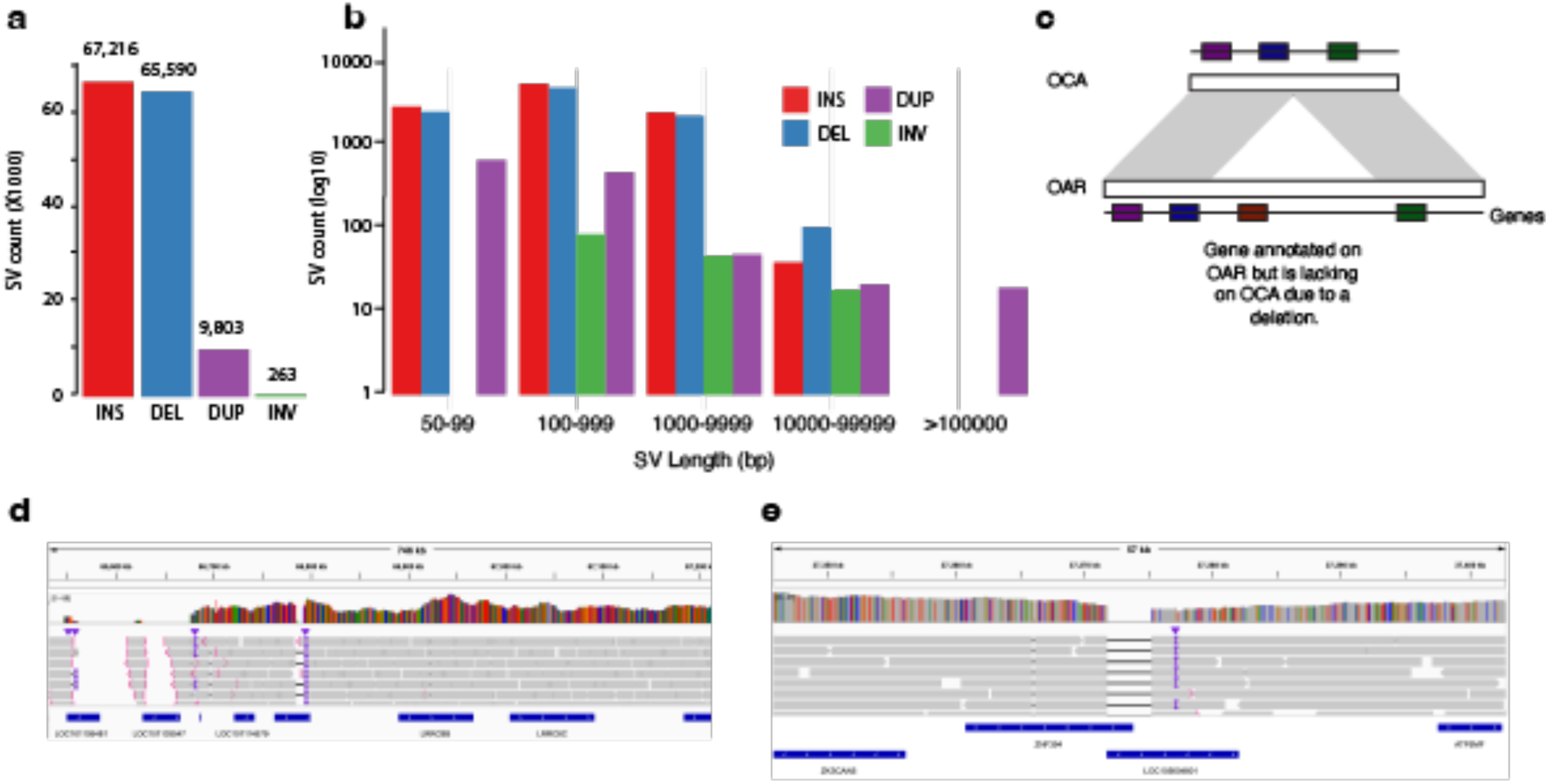
Variants analysis revealing missing immune-related genes on the Bighorn-T2T but present on the domestic sheep. (a) Number of SVs per SV class between bighorn and the domestic sheep as reference (b) The size distribution of the SVs per class showing there are more large SVs than short indels. (c) Sketch diagram illustrating genes missing due to a deletion on the Bighorn-T2T genome (OCA) relative to the domestic sheep genome (OAR). IGV visualization of some of the regions of structural difference between OCA and OAR highlighting two of the genes lacking on Bighorn-T2T within deletions relative to OAR for (d) *GBP5* (*LOC101105481*) and (e) *ZNF501* (*LOC105604801*).

Overall, gene content comparison between bighorn and domestic sheep genomes revealed that 5 protein-coding genes were exclusive to bighorn sheep (Supplementary Data 16). These included *UCMA* (Upper zone of growth plate and cartilage matrix associated), *CFHR4* (Complement factor H-related protein 4-like), *ZNF320* (Zinc finger protein 320-like) and *TTC9C-like* (Tetratricopeptide repeat protein 9C-like) (Supplementary Data 16). Similarly, three protein-coding genes (*LOC101120367* (Guanylate-binding protein 5, *GBP5*), *LOC105604801* (Zinc finger protein 501-like, *ZNF501-like*), and *LOC101115448* (Olfactory receptor 1361-like)) were in the domestic sheep assembly but lacking in Bighorn-T2T (Figure 4c-e, Extended Data Fig.3, Supplementary Data 17).

### Bighorn and domestic sheep adaptive immune systems are characterized by prevalent duplications

To compare the immune loci of bighorn and domestic sheep, five bighorn sheep haplotypes (Bighorn-T2T plus four other haplotypes we assembled from two adult individuals) and 21 domestic sheep haplotypes (obtained from NCBI) were used (Supplementary Data 18).

Annotation of the IG/TR loci revealed that for the V genes cluster, the IGL loci were statistically significantly shorter (P=0.015) in the bighorn sheep haplotypes (0.30– 0.49 Mbp) than the domestic sheep haplotypes (0.39– 0.94 Mbp) (Figure 5a). While the T-receptor TRA/D loci in the bighorn sheep haplotypes (1.29–2.17 Mbp) were also significantly (P=0.006) shorter than the domestic sheep haplotypes (1.43 to 5.26 Mbp), the TRB loci on the bighorn sheep haplotypes (0.39–0.40 Mbp) were slightly but significantly (P=0.006) longer than on the domestic sheep haplotypes (0.26–0.38 Mbp) (Figure 5a). Pairwise alignments of the V gene locus sequences revealed a highly repetitive organization that suggests duplications within corresponding regions in both species (Figure 5b, Extended Data Fig.4). Interestingly, TRA/D loci have the most complex organization compared to other IG/TR loci, and the highest variance in lengths, which contrasts with previous findings^30^ that IG loci are more variable than TR loci. The number of in-frame V genes without stop codons (further referred to as productive) are variable across both species and haplotypes within a species (Figure 5c) and are positively correlated with the lengths of the respective V gene loci within all the locus types except for TRB (Figure 5d, Extended Data Fig.5). Analysis of similarities of V genes within a subject revealed that the average percentage identities of the V genes were significantly higher at the IGK (P=0.002) and TRA/D (P=0.006) loci of the domestic sheep than the bighorn sheep. The same pattern was observed (albeit non-significant) for IGL and TRG loci (Extended Data Fig.5). These observations indicate a higher level of gene duplication in the domestic sheep haplotypes (Figure 5e). The average percent identities of V genes across pairs of the bighorn sheep haplotypes were higher compared to percent identities of V genes computed across pairs of domestic sheep haplotypes and cross-species pairs (bighorn/domestic sheep) across all IG/TR loci with p-values below 0.0001 (Figure 5f).

**Figure 5.**
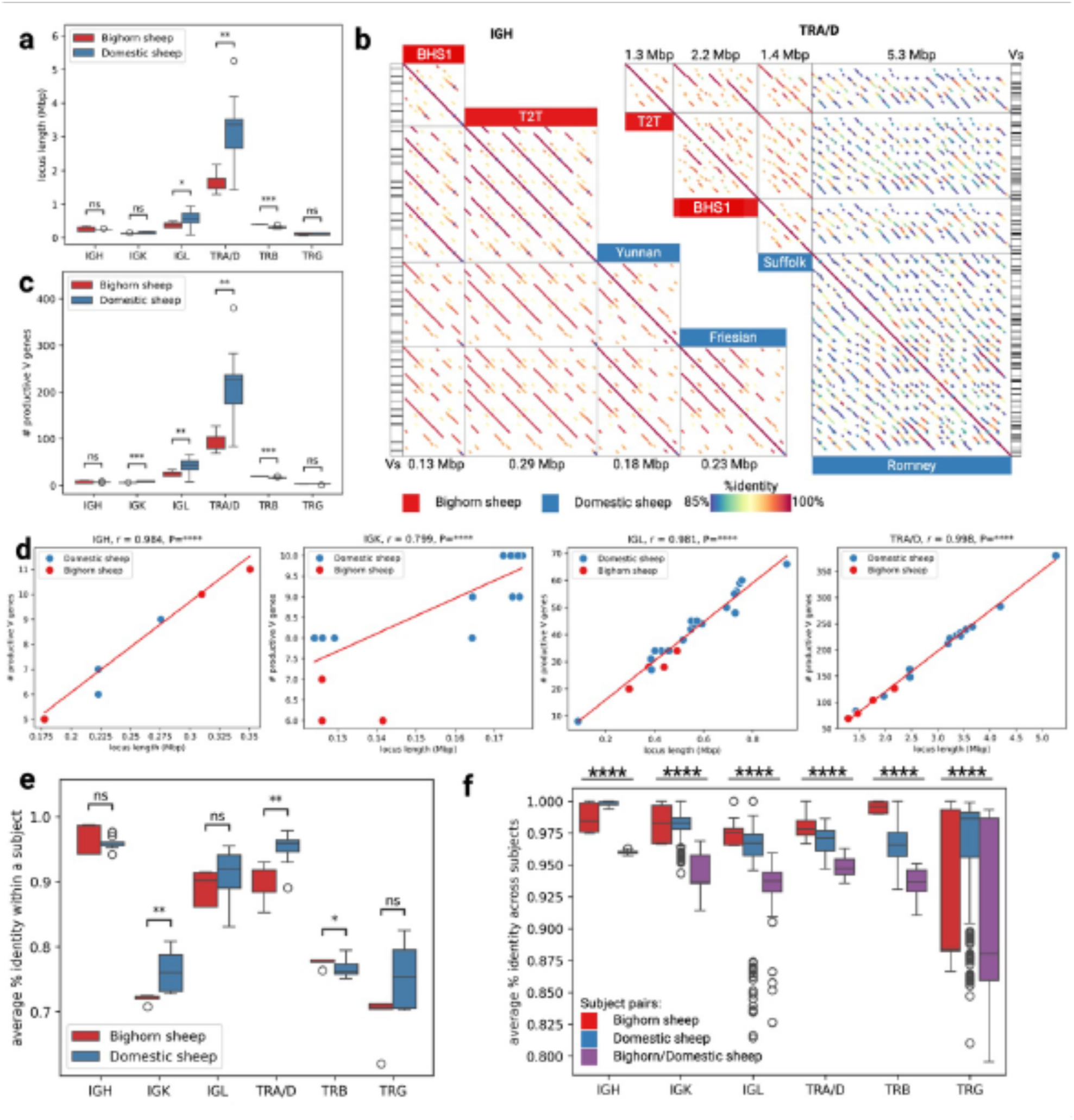
Adaptive immune loci of bighorn and domestic sheep. (a) Distribution of lengths of IG and TR loci in five bighorn sheep and 21 domestic sheep haplotypes indicating that IGL, TRA/D and TRB were statistically significantly different while IGH, IGK and TRG loci were not statistically significantly different (b) Dot plots showing pairwise alignments of four V clusters (from left to right): the shortest bighorn sheep V gene locus, the longest bighorn sheep V locus, the shortest domestic sheep V gene locus, and the longest domestic sheep V gene locus for IGH and TRA/D. Names of the subjects are shown on the corresponding column. Alignments are colored according to their percent identities: from blue (85% and below) to red (100%). Positions of V genes (Vs) are shown on the side. (c) Distribution of counts of in-frame V genes without stop codons collected across IG/TR loci. (d) Correlations between lengths of V gene cluster and counts of in-frame V genes without stop codons for IGH, IGK, IGL, and TRA/D loci, respectively. (e) Distribution of average percent identities of V genes within subjects of the bighorn sheep and domestic sheep across five types of IG/TR loci. (f) Distribution of average percent identities of V genes across all pairs of bighorn and domestic sheep haplotypes across five types of IG/TR loci.

### Innate immune system genes differ between bighorn and domestic sheep

The Leukocyte Receptor Complex (LRC) is a cluster of genes which encode a diverse set of Immunoglobulin-like (IG-like) receptors that control immune cell responsiveness. The LRC in bovids encodes three multi-gene subgroups: the leukocyte IG-like receptors (LILR), killer cell IG-like receptors (KIR), which are expressed on Natural Killer (NK) cells and some T cells, and a third group of yet-to-be-characterized novel IG-like receptors^31^. The LILR and the novel IG-like receptor gene content appeared not to be variable in the bighorn sheep haplotypes, while all three of these gene groups (LILR, KIR and novel IG-like receptor) were expanded and varied considerably in gene content between haplotypes in domestic sheep, including duplications of the bovid-specific *FCG2R* gene (Figure 6A, Supplementary Data 18).

**Figure 6.**
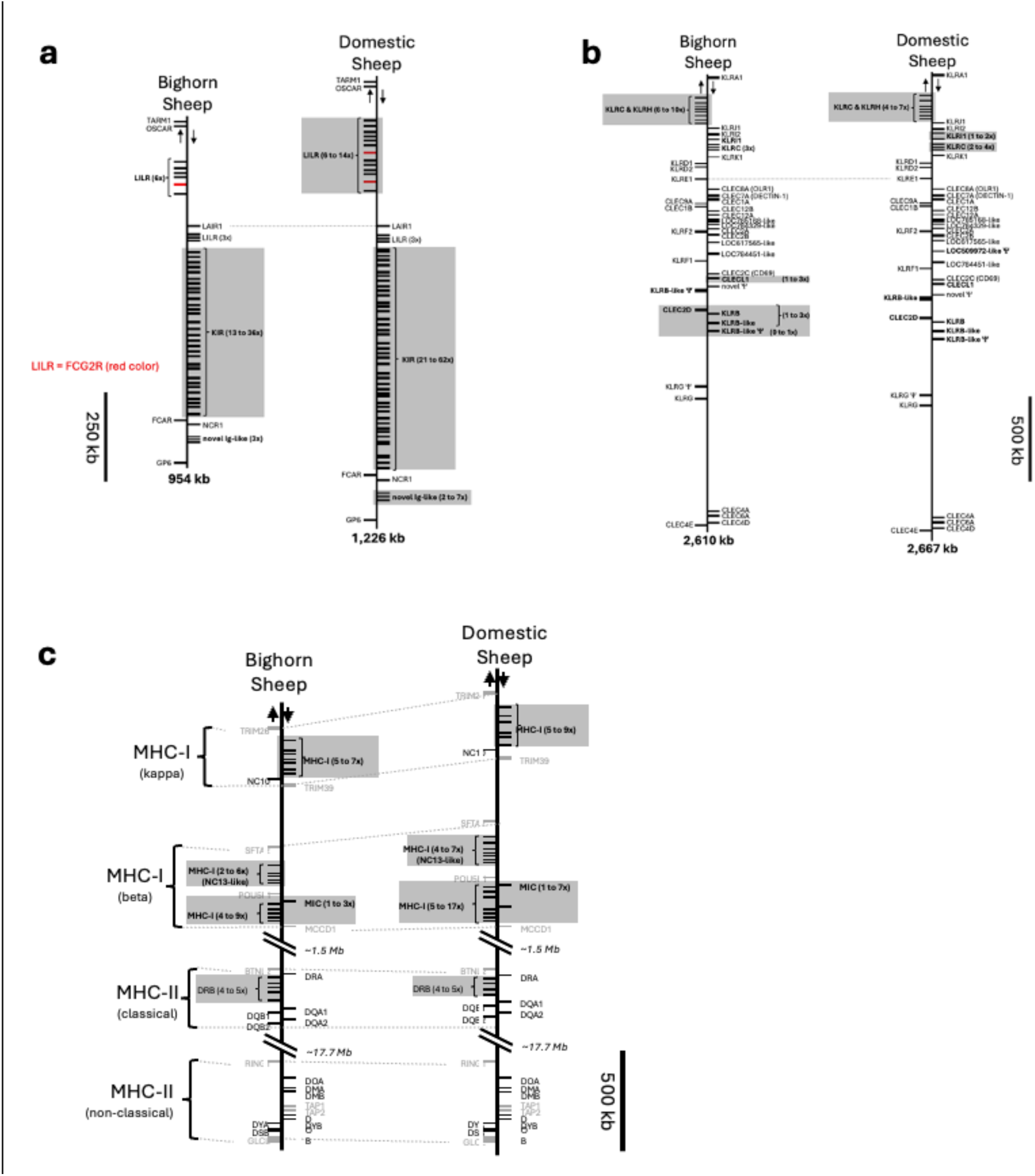
**Organization of the LRC, NKC and MHC immune loci in bighorn sheep compared to the domestic sheep**. For all panels, arrows at the top indicate direction of transcription for genes represented on either side of the vertical backbone. Loci which differ in content between the two species are shown in bold. Paralogous groups of genes which vary in number between haplotypes within a species are shaded in gray, and the range of the number of genes present across the assemblies is indicated. (a) Organization of the LRC in bighorn and domestic sheep. Only IG-like genes are represented. The bighorn and domestic sheep schematics are aligned at LAIR1 (dashed line). The bovid-specific 2-domain LILR gene, FCG2R, is indicated in red. (b) Organization of the NKC in bighorn and domestic sheep. Only C-type lectin genes are represented. The bighorn and domestic sheep schematics are aligned at *KLRE1* (dashed line). (c) Organization of the MHC in bighorn and domestic sheep. MHC-like genes are represented as black bars and conserved non-MHC genes are indicated in gray and serve as positional markers.

The KIR genes are highly expanded and variable in both bighorn (13-36 genes) and domestic sheep (21-62 KIR genes) (Supplementary Data 18). Within the NKC, the *KLRC* and *KLRH* genes, which are flanked by *KLRA* and *KLRJ* genes, showed signs of expansion in bighorn (6 - 10 copies) compared to domestic sheep (4 – 7 copies) (Figure 6b, Supplementary Data 18). In some domestic sheep haplotypes however, duplications of inhibitory *KLRI1* and *KLRC* genes flanked by *KLRI1* and *KLRK* was observed. Furthermore, both functional and non-functional orthologs of the uncharacterized C-type lectin *LOC100138381* (killer cell lectin-like receptor subfamily B member 1) were identified in both bighorn sheep (*LOC138437836*, *LOC138438337*) and domestic sheep. In bighorn sheep, the non-functional copy of this gene, along with *CLEC2D*, *KLRB*, *CLECL1*, and a novel C-type lectin domain-containing pseudogene, was duplicated up to three times on three of the five bighorn sheep haplotypes (Supplementary Data 18).

The MHC-I genes play a major role in immune response by presenting antigen molecules for recognition by TRs. The gene content and organization of the MHC-I Kappa between the bighorn sheep haplotypes (5 – 7 genes) and the domestic sheep haplotypes (5 – 9 genes) (Figure 6c, Supplementary Data 18) were comparable. In contrast, on the Beta block, the non-classical MHC-I genes and the MHC-I-like (MIC) genes exhibited high variability between the bighorn (4 – 9 MHC-I genes and 1 – 3 MIC genes) and the domestic sheep haplotypes (5 – 17 MHC-I and 1 – 7 MIC genes) (Figure 6c, Supplementary Data 18). In addition, a notable annotation in the beta block is the *NC13-like* gene (previously *P2*^32^) where 2 – 5 copies were annotated in the bighorn sheep haplotypes compared to 4 – 7 copies on the domestic sheep haplotypes (Supplementary Data 18). The MHC-II on the other hand did not show variability in the gene content between the bighorn and the domestic sheep haplotypes except for an additional *DRB* pseudogene which was occasionally observed in the haplotypes of both species (Figure 6c).

### Utility of Bighorn-T2T in pathogen (M. ovipneumoniae) susceptibility studies

Martin *et al.*^33^ endeavored to determine how genomic composition of bighorn sheep affects the persistent carriage of *M. ovipneumoniae* without the aid of a quality reference assembly. Their study examined a bighorn sheep population for which longitudinal disease status data (including sinus tumor presence) were available using RAD-Seq with the domestic sheep Oar_v4.0^34^ assembly as reference. A total of 10,605 SNP loci from 25 individuals were used in the analysis to identify two SNPs of interest and seven cis-candidate genes located on OAR1 and OAR7^35^.

When reanalyzing this data using Bighorn-T2T as reference, the RAD-seq reads aligned better to the Bighorn-T2T (Figure 7a) (avg. 75.16% properly paired reads) in comparison to the older OAR reference genome (GCA_000298735.2, Oar_v4.0) used in the original study^33^ (avg. 62.60%) as well as the current (avg. 70.80%) OAR reference genome (GCA_016772045.2, ARS-UI_Ramb_v3.0) (Figure 7a). We identified five SNPs of interest from 54,641 loci (Figure 7c, Table 2) which differed from the two SNPs originally identified^33^. Four of the five SNPs were in intergenic regions, and one was located within Calpain 2 (*CAPN2*) on OCA12 (Table 2). The SNP in *CAPN2* (effect allele = A, reference allele = G) exhibited a strong association with *M. ovipneumoniae* carrier status (Beta = 0.774, p = 3.19E-05). Genotype distribution revealed about 85% GG and 15% GA individuals, with no AA observed (Figure 7b). This distribution corresponds with a higher effect allele (A) frequency in cases (0.417) compared to controls (0.053), indicating a direction towards increased susceptibility. A total of 22 protein-coding genes (four with immune-related functions) were located within 100kb of these SNPs (Table 2).

**Figure 7.**
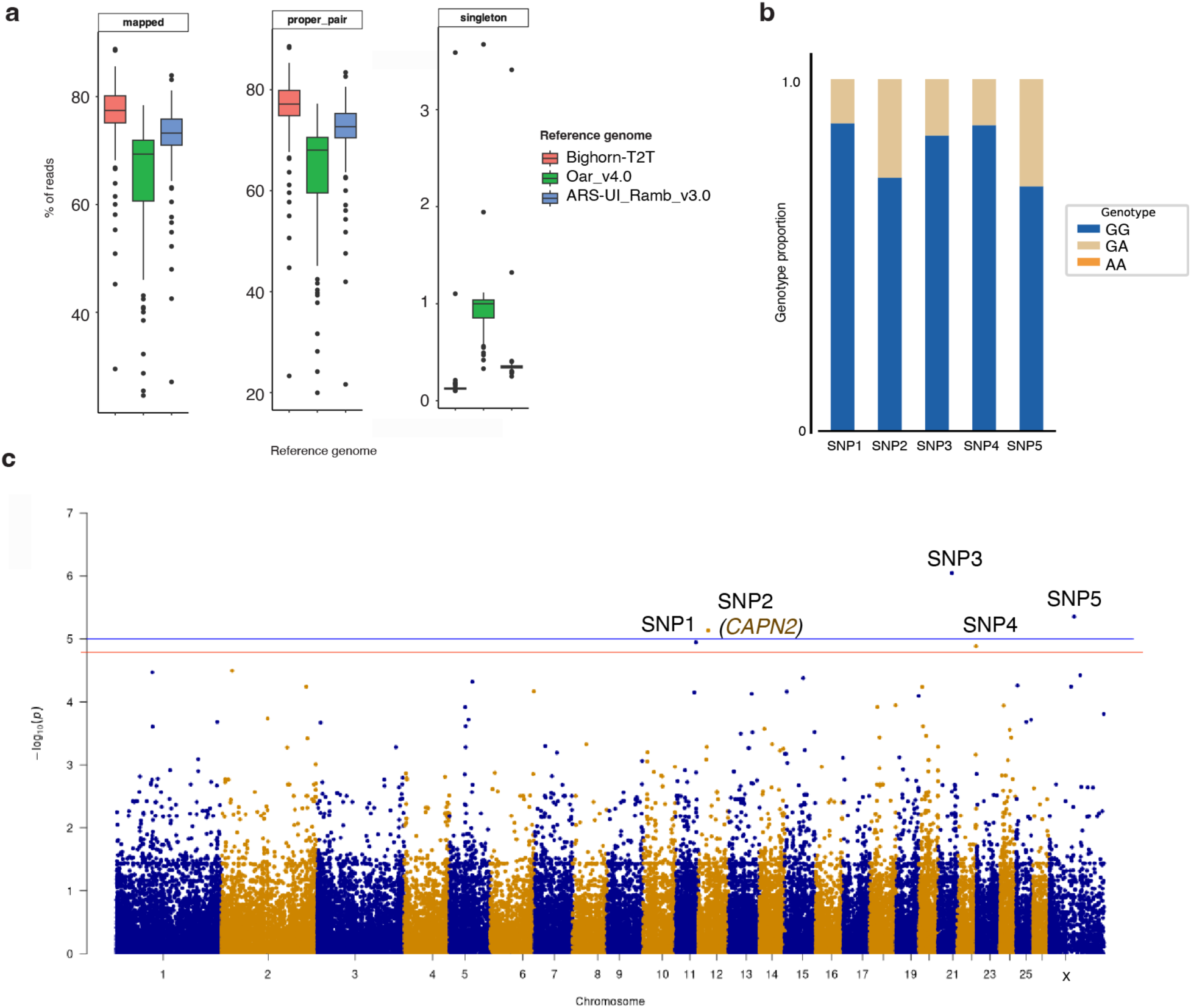
**Reanalysis of *M. ovipneumoniae* chronic carrier and sinus tumor status in bighorn sheep**. (a) Mapping rates of the RAD-seq sequence obtained from Martin *et al*.^35^ to the Bighorn-T2T and the domestic sheep Oar_v4 and Ramb_v3.0 assemblies. The bighorn sheep assembly exhibited the highest mapping rates of the three assemblies. (b) Genotype distribution of the SNPs of interest for the 25 individuals with longitudinal disease carriage data that were used in the study. (C) Manhattan plot of the reanalysis using Bighorn-T2T as the reference highlighting identified loci of interest. The genome-wide significance level is defined with the blue line and the red line is the p-value at the applied 0.15 FDR.

**Table 2:** Identified SNPs of interest from reanalysis of *M. ovipneumoniae* carrier status^33^ in bighorn sheep using Bighorn-T2T as a reference genome. The variant effect size which is reported as the Beta values; positive values are associated with susceptibility while negative values are associated with resistance. The alternate allele frequency of a SNP for case and controls are Case AF and Cotrol AF, respectively.

| SNP | CHR | POS (bp) | REF | ALT | Beta** | Case AF | Control AF | Effect | Likelihood ratio test p-value | Genes within 100kb flanking SNP locus |
| --- | --- | --- | --- | --- | --- | --- | --- | --- | --- | --- |
| SNP1 | 11 | 64,127,679 | A | C | 0.940 | 0.300 | 0.000 | ↑(susceptibility) | 7.58E-05 | *LLGL2* (LLGL scribble cell polarity complex component 2), *TSEN54* (tRNA splicing endonuclease subunit 54), *CASKIN2* (CASK interacting protein 2), *TMEM94* (transmembrane protein 94), SNP1, *GRB2* (growth factor receptor bound protein 2) |
| SNP2 | 12 | 37,621,644 | G | A | 0.774 | 0.417 | 0.053 | ↑(susceptibility) | 3.19E-05 | *CAPN8* (calpain 8), *RPL23A*, (ribosomal protein L23a), ***CAPN2* (calpain 2)\***, *TP53BP2* (tumor protein p53 binding protein 2) |
| SNP3 | 21 | 53,140,775 | A | G | 0.904 | 0.333 | 0.000 | ↑(susceptibility) | 5.00E-06 | *CD6* (CD6 molecule)*, SNP3, *CD5* (CD5 molecule)*, *VPS37C* (vacuolar protein sorting 37C), (*PAG4*, pregnancy-associated glycoprotein 4) |
| SNP4 | 22 | 66,312,702 | G | A | 0.936 | 0.300 | 0.000 | ↑(susceptibility) | 8.31E-05 | *PWWP2B* (PWWP domain containing 2B), *LOC138427599*, SNP4, *INPP5A* (inositol polyphosphate-5-phosphatase A) |
| SNP5 | X | 65,177,879 | T | C | 0.714 | 0.500 | 0.056 | ↑(susceptibility) | 2.53E-05 | *SNX12* (sorting nexin 12), *FOXO4* (forkhead box O4), *CXHXorf65* (chromosome X open reading frame 65)*, *IL2RG* (interleukin 2 receptor, gamma chain)*, *MED12* *, SNP5, *NLGN3* (neuroligin 3), *GJB1* (gap junction protein beta 1), *ZMYM3* (zinc finger MYM-type containing 3) |

Gene in bold denotes the SNP is within that gene, \*Genes with immune function

## DISCUSSION

Bighorn-T2T was produced by leveraging a combination of technologies involving short reads, HiFi, ultra-long and HiC sequence data which has been successfully employed in other mammalian T2T assemblies^15,17^. In agreement with previous reports in other mammalian species, more than half of the bighorn sheep genome is repetitive. Bighorn-T2T adds NARs from the telomeres, centromeres, rDNA arrays, segmental duplications, and other DNA amounting to over 400Mb (14.28% of the genome) of previously unobserved genomic features in Bighorn-v1. Newly assembled genes which were not observed in Bighorn-v1, including genes with immune-related function can now be incorporated into bighorn sheep genomic studies and other genetic resource development.

The NORs which are responsible for ribosome biogenesis^36^ and nucleolus formation^37^ were newly assembled in Bighorn-T2T. Contrary to the expected number of arrays on the domestic sheep^26^, no NOR was observed on the distal p-arm of OCA2 where it was expected in addition to arrays on OCA1, OCA3, OCA4, and OCA25. We hypothesize that this lack of an rDNA array on OCA2 suggests polymorphism of rDNA in bighorn sheep as observed in primate species^15^.

Confirmation of this polymorphism and the extent of the variation in ruminants also remain to be tested on multiple individuals across species^38^.

Structural genomic differences between bighorn and domestic sheep, which also involved genes, underscores the observed dissimilarity, and could potentially contribute to the genetic underpinnings of respiratory disease pathogenesis and clinical phenotype distinction between the two species. The shorter length of OCAY compared to OARY was due to fewer ampliconic gene families on the MSY region on OCAY, suggesting loss of some of these gene families (mainly the bovine-specific^39^ *ZNF280B-like* genes and the ancestral mammalian *TSPY* already pseudogenized on the domestic sheep) in bighorn sheep since the split from domestic sheep about 5.3MYA^1^. *UCMA*, one of the genes in bighorn but not in domestic sheep encodes chondrocyte-specific, highly charged proteins that are abundantly expressed during the early stages of chondrogenesis^40^. Despite the implied role in calcification and ossification, mice lacking the encoded protein do not display significant defects in the skeletal development^41^. *CTPS1*, another of the genes within a region of variation between the two species was predicted to have a high impact on exons 5-8 of 20 in the bighorn sheep. A loss of function of *CTPS1* by splicing out exon 18 has been reported to result in life-threatening immunodeficiency in humans^42^, impairing the ability of T and B cells to proliferate upon antigen receptor-mediated activation. In addition, two of the three genes identified in the domestic sheep and lacking in the bighorn sheep genome, *GPB5* (Guanylate-binding protein 5) and *ZNF501-like* (Zinc finger protein 501-like), are associated with the innate immune system. Perhaps the most compelling discovery of Bighorn-T2T is the lack of *GBP5*. *GBP5* promotes NLRP3 (NOD-like receptor protein 3) inflammasome assembly, an inflammasome known to play a critical role in the innate immune response against parasitic, fungal, bacterial, and viral infections ^43,44^. Consequently, the lack of *GBP5* can be expected to parlay into increased risk of pathogen susceptibility, including pathogens associated with respiratory diseases. Mice deficient for NLRP3 expression have decreased immune cell activation and delayed bacterial clearance during acute infection with *M. pneumoniae*^45^. In humans, reduced expression of *GBP5* has been linked to weaker immune response and greater susceptibility to viral infection^46^*. GBP5* anti-viral activity has been linked to its sub-cellular localization to the trans-Golgi network in humans^47,48^ and in bats^49^. *ZNF501* is thought to be involved in Golgi organization and is essential for its structural integrity and thereby protein packaging functionality^50^. Although localization of pathogens to the Golgi network has been implicated in promoting pathogen proliferation^51,52^, the Golgi has also been reported to be required to activate a STING-dependent signaling for the type-1 IFN response^53^. This study establishes the first reported hypothesis that the lack of these two critical immune response genes in bighorn sheep could be a major determinant of the differential response of bighorn sheep to *M. ovipneumoniae* infection.

Furthermore, in the innate immune system, the expansion and variability in the copies of KIR genes in both bighorn and domestic sheep agrees with reports in bovids, which reported were the only known species to feature expanded gene content of both LILR and KIR beyond primates^54–56^. KIR and LILR genes are critical in maintaining the balance between self-tolerance towards normal cells and inflammation regulation^57,58^. The variability of LILR on domestic sheep potentially confers the ability to diversify functionality^59^ and widen pathogen range through increased receptor variants^60^.

The *KLRC* and *KLRH* genes within the NKC of bighorn sheep were observed to be expanded relative to domestic sheep. *KLRC/KLRD* heterodimeric receptors in human and mice recognize nonclassical MHC-I ligands while *KLRI1* is an inhibitory NKC receptor which heterodimerizes with *KLRE* in rodents and interacts presumably with an unidentified MHC-I ligand^61^. The expansion of the *CLEC2D* and *KLRB* in bighorn sheep relative to domestic sheep is notable due to their roles in innate immune response. *CLEC2D* forms homodimers and is ubiquitously expressed across various cell types, where it can act as a sensor for cell death and modulate immune responses via recognition by inhibitory *KLRB* receptors (also known as *NKR-P1*) present on NK cells^62^. The duplicated bighorn sheep *CLEC2D* paralogs are all identical in amino acid sequence except for a single residue in the predicted transmembrane region of one paralog (Cys to Tyr). In contrast, the duplicated *KLRB* paralogs differ by as much as 28 residues (88 percent identity). Paralogs could evolve tissue-specific or function-specific roles. Taken together, the observed similarity and the difference in the paralogs of *CLEC2D* and *KLRB-like* receptors, respectively, suggests an expanded role of *KLRB*-like receptors in bighorn compared to domestic sheep.

In the variable region of the adaptive immune receptors comprising the V gene cluster, only the IGL loci showed a difference between the species while the IGH and IGK were comparable.

Generally, the IG loci were much shorter in bighorn than in the domestic sheep haplotypes. The number of productive V genes exhibited a direct correlation with the length of a locus, except for TRB locus which was longer in bighorn sheep. These shorter loci on the bighorn sheep reduces the size of receptor repertoires owing to combinatorial restriction of the genes^11,12^. As anticipated, the gene content exhibited high variability for the different gene types. The V genes were found to have lower sequence identity between the copies at the bighorn sheep IGK, IGL and TRA/D loci relative to domestic sheep, which is indicative of a lower level of gene duplication in bighorn compared to domestic sheep. However, higher inter-species haplotype variation is expected in domestic sheep potentially due to multiple breed selection pressure, in comparison to the wild bighorn sheep^8^.

The average percent identities within species were higher compared to cross-species pairs across all IG/TR loci suggesting that independent evolution of the bighorn and domestic sheep lineages has resulted in species-specific mutations of adaptive immune genes. However, the results also indicate that IG and TR loci likely evolved at different rates. Although there was a lack of multiplicity in the IG loci between the two species, the high level of multiplicity observed in TRs is contrary to recently reported findings in primates^15^. Generally, multiplicity of genes in the IG and TR loci are beneficial as it presents the species with increased variability in recognizing and responding to invading pathogens.

The significance of Bighorn-T2T genome as a resource for bighorn sheep genomic studies has been further underscored with the different outcome of re-analysis of a *M. ovipneumoniae* pathogen susceptibility study in bighorn sheep. Contrary to two SNPs and seven candidate genes identified in the original study using the domestic sheep as the reference, we identified five SNPs, including one located in Calpain 2 (*CAPN2*) which has been implicated in a range of physiological activities of immune cells including TR activation under pathological conditions^63–65^. Four more genes out of the total 25 genes within 100kb of the identified SNPs of interest have immune-related functions. These results point to the value of the Bighorn-T2T as a genomic resource.

### Limitations of the study

The unavailability of immune repertoire sequencing (Rep-Seq)^66^ data on bighorn and domestic sheep for this study prevented further investigation of the dynamics of adaptive immunity in the two species. Given that T-and B-cell receptors are generated through V(D)J recombination sequencing, the Rep-Seq data would have enabled analyses that provide further insights into antigen recognition and immune response in the two breeds. Another limitation of this study is the likely data bias in the comparison of the IG/TCR loci of the two species since there were more domestic sheep haplotypes (21 haplotypes) than bighorn sheep haplotypes (5 haplotypes) used in the analysis. The higher average percent identities of V genes recorded in bighorn may have been due to the imbalance in the number of samples. Finally, the sample size of 25 used in the bighorn sheep pathogen carriage status study^33^ that we re-analyzed was low for the genome-wide association study (GWAS). A larger sample size would have increased the statistical power of the GWAS and reduced the risk of false negatives thereby increasing confidence in the results.

## SUMMARY

The first Bighorn sheep T2T genome assembly presented here has resolved the full architecture of the immune loci and revealed a critical immune-response gene, *GBP5,* and an immune response-mediating gene, *ZNF501,* present in domestic sheep but absent in bighorn sheep.

Specifically, the absence of *GBP5* offers a strong hypothesis for the increased susceptibility of bighorn sheep to respiratory pathogens such as *M. ovipneumoniae*. Bighorn-T2T sheep reference has identified immune-related candidate genes linked to pathogen susceptibility, including *CAPN2*. These findings establish Bighorn-T2T as a critical genomic resource for understanding immune system differences, disease vulnerability, evolution, and the wider genetic differences that shape phenotypic divergence between bighorn and domestic sheep.

## METHODS

### Ethics statement

Animal care and handling for Polypay sheep was under the auspices of the Washington State University (WSU) Institutional Animal Care and Use Committed (IACUC) ASAF 6853. The additional Bighorn sheep samples were collected under WSU IACUC ASAF 4885. The University of Idaho held the project oversight IACUC-2020-58.

### Interspecies fetuses

Purebred unrelated multiparous Polypay ewes (5 and 3 years old), known for increased prolificacy, were artificially bred with bighorn sheep ram semen from two different bighorn sheep rams. Tissues were collected and flash frozen from ewes and fetuses within 40 minutes of death and blood samples were collected just before or after death (Extended Data Fig.1, Supplementary Methods).

### Nucleic acid extraction and NGS library preparation and sequencing

High-molecular-weight (HMW) DNA was isolated from tissues utilizing a HMW Phenol:Chloroform protocol. Details of the library preparation and sequencing can be found in Supplemental Methods.

### Pushing the bighorn sheep assembly to T2T

Verkko (v2.0) produced 18 autosomes and the Y chromosome as telomere-to-telomere (T2T) chromosomes comprising 15 T2T contigs and 4 T2T scaffolds (Supplementary Data 3). Telomeric contigs were anchored on the chromosomes by aligning the ONT UL reads to the assembly graph (See Supplemental Methods).

### Assembly Polishing

The manually curated assembly was polished using the T2T-Polish pipeline^67^ based on Merfin and Racon. The HiFi reads were aligned using Winnowmap2 while the high-quality hybrid-kmer database produced with Merqury^21^ and Meryl were used for the polishing (see details in supplementary methods).

### Assembly completeness evaluation

Compleasm^20^ was run on the assembly with default parameters to evaluate the number of BUSCOs retrievable from the assembly using the *Cetartiodactyla_odb10* database as a proxy for the assembly completeness evaluation.

### Bovine and ovine satellites annotation

The blast output was filtered for alignments with a minimum of 1E-3 e-value and 80% coverage between the query and the target sequence. The Bovine satellite alignments were however filtered with 80% coverage between the query and the target due to the higher divergence between the satellites and the targets on the bighorn sheep genome.

### Repeats annotation

RepeatMasker^68^ was used to annotate and mask the bighorn sheep assembly while RepeatModeler^69^ was employed for de novo repeats discovery. The new repeat *consensi* sequence from RepeatModeler were subsequently run on TEtrimmer^70^ to refine the boundaries and generate new consensus sequences.

### Analysis of structural variation with domestic sheep

The PacBio HiFi reads of the Bighorn_x_Polypay F1 cross were trio-binned with Canu^71^ using the parental Illumina data as markers. The binned HiFi reads from the bighorn sheep haplotype were then aligned to the domestic sheep reference genome with the PacBio Minimap2^72^-based Pbmm2 (https://github.com/PacificBiosciences/pbmm2). Structural variants were called against the domestic sheep reference genome with Minimap2 and Paftools.

### Segmental duplications across the genome

Segmental duplications (seg dups) across the genome were analyzed with Biser^73^ using soft-masked version of the assembly with RepeatMasker^68^. The results were filtered for at least 90% identity between alignment blocks of at least 1kbp length. Duplicate records were filtered with in-house scripts and the inter-and intra-chromosomal events were segregated. The seg dups loci were further filtered to exclude loci with at least 70% coverage overlap with the consolidated repeats annotation.

### Immune loci comparison

We investigated the interspecific differences between bighorn and domestic sheep using multiple haplotypes due to the known polymorphic nature of the immune loci within and between species. The four additional bighorn sheep haplotypes generated from HiFi reads of two pure-bred animals and 21 domestic sheep haplotypes were used to elucidate the immune loci of the two species. The findings at the adaptive and the innate immune loci between the two species are highlighted as follows.

### IG/TR loci V genes similarity analysis

To analyze V gene similarity, all pairs of subjects (s1, s2) were analyzed. Then, for each V gene from s1, the closest V gene from s2 was found and the corresponding percent identity was computed. For each pair (s1, s2), the computed percent identities were collected from all V genes and the average percent identity was computed. In case s1 and s2 represent the same subject, the closest V genes within the same locus were found, and the average percent identity was referred to as the average subject % identity.

### Manual annotation and curation of innate immune-related gene complexes

The MHC class I and class II genes, C-type lectins encoded within the NKC, and Ig-like genes encoded within the LRC were initially annotated using parsed BLASTN searches with manually annotated and/or curated sequences available for sheep, goat, and cattle^56,74–76^. These initial annotations were then manually refined using Artemis^77^ and gene content compared. Manually annotated MHC alleles from the paternal bighorn sheep haplotype were submitted to the sheep section of the ImmunoPolymorphism Database (IPD-MHC)^76^ for curation as prototypic alleles for the species.

### Methylation analysis

The methylated CpGs were analyzed from the ONT reads using the prescribed analysis pipeline. The fastq reads files were extracted from the *modbams* with the *MM, ML* tags retained and aligned to the bighorn sheep assembly using Winnowmap^78^. The alignment file was parsed with modbam2bed, and the output bed file was processed into bigwig file using the UCSC *bedGraphToBigWig* module to enable visualization of the tracks on IGV.

### Centromeric region self dotplots

The self-identity dotplots of the centromeric region of all the chromosomes were made with Moddotplot (https://doi.org/10.1101/2024.04.15.589623) (see supplementary methods for details).

## Data Availability

The Bighorn-T2T genome reported in this study has been deposited in GenBank under the accession GCA_042477335.2 and can be accessed at the URL https://www.ncbi.nlm.nih.gov/datasets/genome/GCF_042477335.2/

## Acknowledgements

This research was supported by the intramural research program of the U.S. Department of Agriculture, National Institute of Food and Agriculture, award numbers 2021-67016-33416, 2023-67015-39000, and National Animal Genome Research Program - Multi State Hatch Project No. IDA01782 and IDA01566.

The results reported here were made possible with computational resources provided by the University of Idaho Research Computing and Data Services (RCDS), FALCON (https://doi.org/10.7923/falcon.id), the NIH HPC Biowulf cluster (https://hpc.nih.gov) and the USDA shared compute clusters Ceres and Atlas as part of the ARS SCINet initiative. This work was also supported, in part, by the Intramural Research Program of the National Human Genome Research Institute, National Institutes of Health. The authors wish to thank Emma Karel-Ward and Kirstin Stena Erickson for animal management and care; Kaneesha Hemmerling, Emma Karel-Ward, Kirstin Stena Erickson, Darren Schneider, Maria Herndon, Gabi Becker, Katie Shira, Gwenyth Potter, Samantha Amey-Gonzalez, Morgan Burke, Gabrielle Condino, Kimberly Davenport, Codie Durfee, Kelly Liebers, Mieko Schwartzmiller, Carolyn Thornberry, Patricia Villamediana, and Breckan Waite for sample collection. Any mention of trade names or commercial products is solely for the purpose of providing specific information and does not imply recommendation or endorsement by the U.S. Department of Agriculture. The USDA is an equal opportunity provider and employer.

## Extended Data

**Extended Data Fig. 1.**
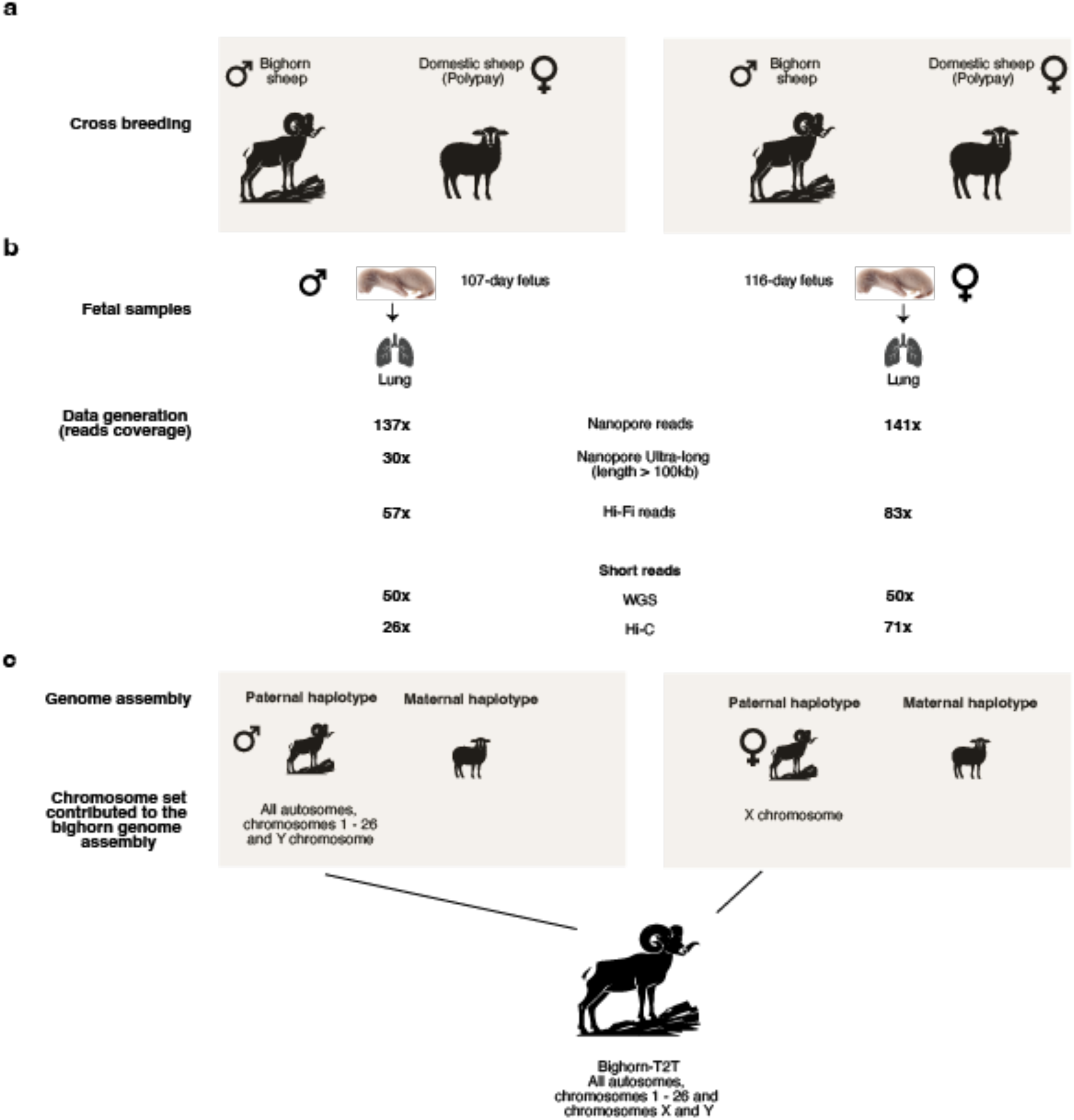
sample collection, sequencing, and genome assembly of the bighorn sheep. (a) Cross breeding between two different Bighorn sheep rams and two unrlated Polypay domestic sheep dams. (b) Fetus age, tissue samples collected and the depth of sequencing produced from The different technologies employed. (c) The set of chromosomes obtained from the samples lo make up The complete Bighom T2T assembly.

**Extended Data Fig. 2.**
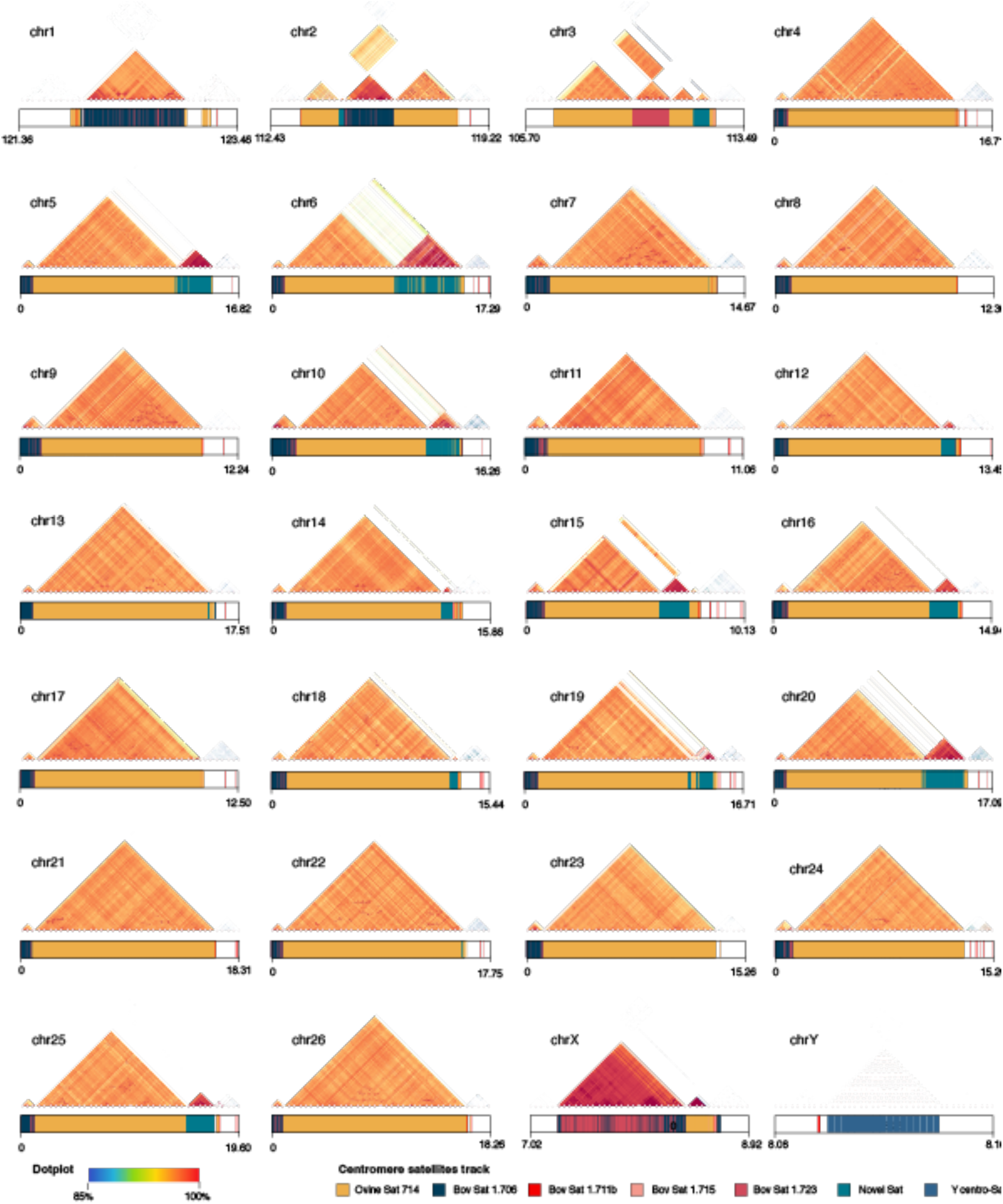
Content and organization of bighorn sheep centromeres. The dotplot of the centromeres showing sequence identity heatmap with the content and organization of the ovine, bovine and the newly annotated centromeric satellites shown in the tracks below (sizes not to scale but the coordinates are in Mbp). The Ovine Sat 1.714 is the most abundant at the centromeres of all the autosomes, except on chromosome 1whιch had the shortest length. The newly annotated centromeric satellite, where present, exhtxted higher sequence identity between the copies, seen on the dotplots.

**Extended Data Fig. 3.**
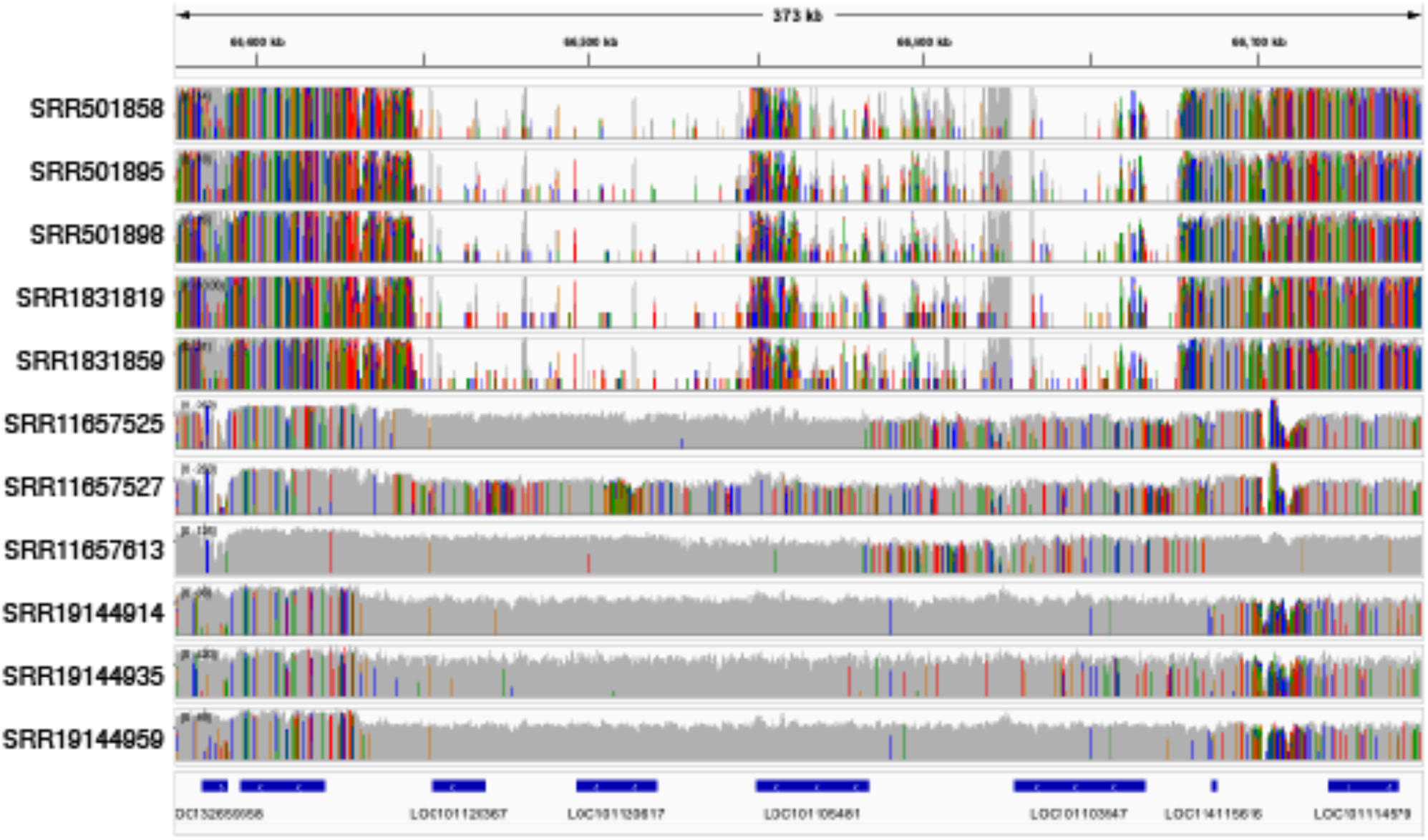
Genes in regions of deletion on the bighorn genome. The whole genome sequence short reads of bighorn (first 5 tracks) and domestic sheep (last 6 tracks) aligned to the domestic sheep reference genome. The alignment of the bighorn reads confirm the deletion of these regions on the bighorn genome. The last track shows the gene annotation of the domestic sheep genome highlighting *GBP5* (*LOC101105481*) in the deleted regions of the genome.

**Extended Data Fig. 4.**
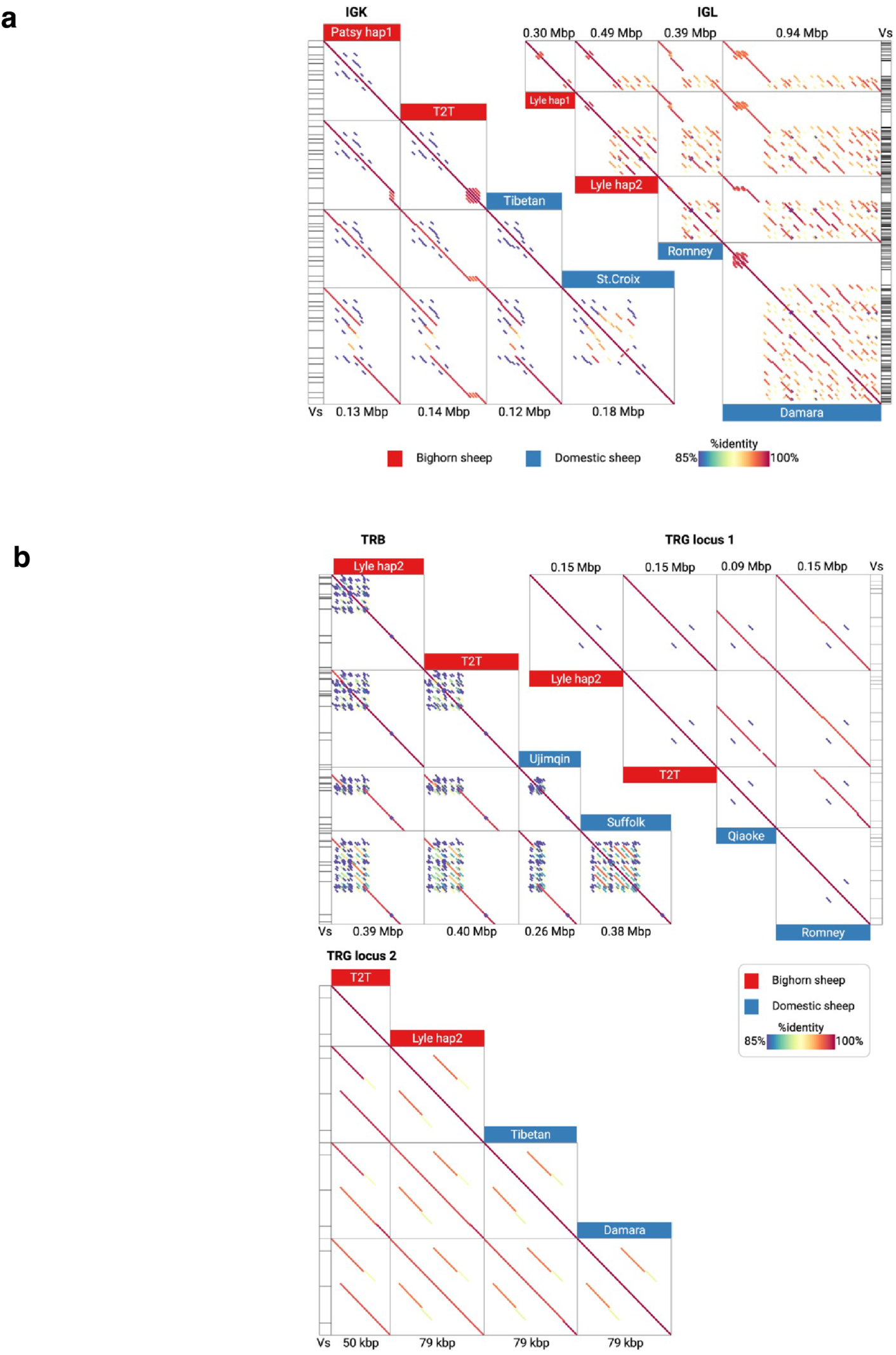
Genomic organization of IG and TR loci on Bighorn-T2T. (a) Genomic organization of IGK and IGL loci in the bighorn sheep and the domestic sheep. Pairwise dot plots of sequences of the shortest and longest V gene region in both species. (b) Genomic organization of TRB and both TRG loci in the bighorn sheep and the domestic sheep. Pairwise dot plots of sequences of the shortest and longest V gene region in both species. The legend is consistent with Fig. 5b.

**Extended Data Fig. 5.**
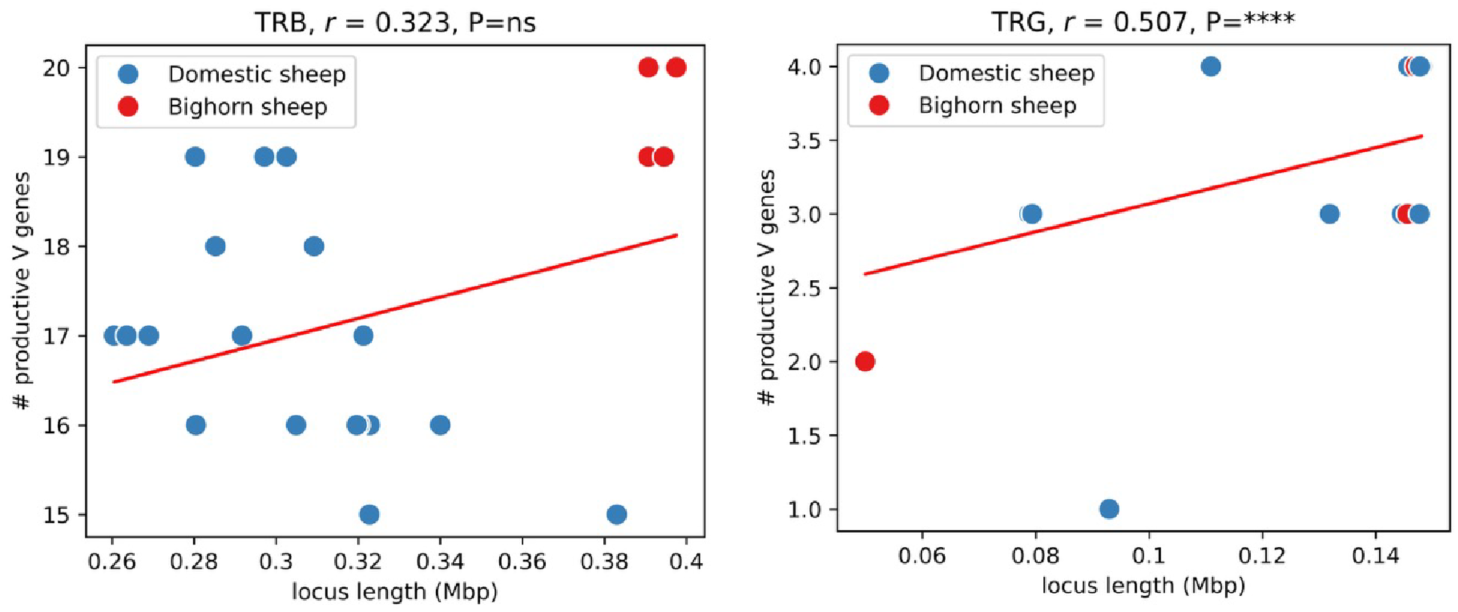
Correlations between the counts of productive V genes and lengths of genomic regions spanning them in TRB (left) and TRG (right) loci. For TRG loci, both proximal regions were used.

