## Supplementary Methods for "Bighorn sheep T2T genome assembly reveals differences in immune genes: a potential cause of high morbidity due to respiratory pathogens"

#### Table of Contents

***1.0 Animal samples ... 2***

**1.1 Interspecies fetuses ... 2**

***2.0 Nucleic acid extraction, library preparation and sequencing ... 2***

**2.1 DNA and RNA extraction ... 2**

**2.2 Library Preparation ... 2**

2.2.1 Pacific Biosciences ... 2

2.2.2 Oxford Nanopore ... 3

2.2.3 Illumina ... 4

***3.0 Estimation of Heterozygosity ... 5***

***4.0 Bighorn genome assembly ... 6***

**4.1 Male F1: All autosomes and chromosome Y ... 6**

4.1.1 Manual assembly curation ... 6

4.1.2 Assembly Polishing ... 14

**4.2 Female F1: Chromosome X ... 15**

***5.0 Validation of nucleolar organizer region (NOR) rDNA array lack on OCA2 ... 16***

***6.0 Repetitive sequence annotation ... 20***

**6.1 Ovine and bovine satellites annotation ... 20**

**6.2 (Peri)centromeric repeats ... 21**

**6.3 RepeatMasker annotation ... 23**

**6.4 De-novo repeats discovery and annotation ... 23**

***7.0 Structural difference analysis ... 28***

**7.1 PBSV variants call ... 28**

**7.2 Minimap2 variant calls ... 28**

**7.3 Variants effect prediction with SNPEff ... 29**

***8.0 Newly assembled genes ... 29***

***9.0 Evaluating the biological significance of a bighorn T2T assembly for population studies ... 31***

***10.0 Self-identity dotplots ... 34***

### 1.0 Animal samples

#### 1.1 Interspecies fetuses

Purebred unrelated multiparous Polypay ewes were artificially bred with Bighorn ram semen from two different Bighorn rams. Ewes were fed to meet or exceed NRC recommendations until sample collection at 107- (male) and 116 (female)-days gestation. The aim was for fetal development similar to the 100 days gestation of domesticated sheep, however there is approximately 35 days difference between domestic and bighorn sheep gestation lengths leading to different gestational ages of the cross-species samples. All tissues were collected, and flash frozen from ewes and fetuses within 40 minutes of death and blood samples were collected just before or after death.

### 2.0 Nucleic acid extraction, library preparation and sequencing

#### 2.1 DNA and RNA extraction

High-molecular-weight (HMW) DNA was isolated from tissues using a HMW Phenol:Chloroform protocol. RNA was isolated from tissues utilizing a TRIzol Reagent and Qiagen RNAeasy protocol.

#### 2.2 Library Preparation

##### 2.2.1 Pacific Biosciences

Pacific Bioscience (PacBio) HiFi libraries were generated for the male and female Bighorn_x_Polypay F1 fetal samples. DNA was sheared using a Diagenode Megaruptor 3 to 20 kb mode size. At all steps, DNA quantity was checked on a Qubit Fluorometer with a dsDNA BR Assay kit, and sizes were examined on a Fragment Analyzer. SMRTbell libraries were then prepared for sequencing according to the manufacturer-prescribed protocol. The libraries were then size selected on the Blue Pippin using the BLF-7510 cassette under the 0.75% DF Marker S1 high-pass 15-20kb protocol to remove fragments below 15kb in size. The selected library fractions were either bound with Sequel II Binding Kit 3.2 (Male F1 Fetus) or the Revio Polymerase Kit (female F1 fetus) and sequenced on Sequel II or Revio instruments (PacBio) with 30 h movie time. Samples were sequenced to a minimum HiFi data amount of 160 Gbp (55x estimated genome coverage).

##### 2.2.2 Oxford Nanopore

Library preparation was done using the Oxford Nanopore Technologies (ONT) Ligation Sequencing Kit LSK-110 (For the male F1 fetus) and the LSK-114 (for the female F1 fetus), followed by ultra-long read sequencing on the PromethION instrument. 10 µg of DNA was cleaned and concentrated by addition of 1 volume of AMPureXP bead solution. After collecting the beads by magnet, the beads were washed 2 times with 0.5 mL 80% ethanol and dried for 1 min. The beads were suspended in 51 µL elution buffer (EB) and DNA eluted at 37°C for 15 min. The eluted DNA was then processed by ligation to the ligation adapter (AMX) as recommended by the manufacturer’s instructions, except increasing the room temperature incubation of the ligation reaction to 60 minutes. Libraries were sequenced in 6 flow cells from two separate template preparations on a PromethION instrument using R9.4.1 flow cells (For the male F1 fetus) and R10.4.1 flow cells (for the female F1 fetus).

##### 2.2.3 Illumina

Parental data of the male F1 fetal cross was obtained from a splenic sample of the Polypay dam. DNA was then processed into short read libraries using the TruSeq DNA PCR-Free Kit as recommended by the manufacturer and sequenced on a NextSeq2000 platform using 2 × 151 base paired end reads. 50x coverage data was generated for the parental data as well as for the F1 fetal cross.

Illumina Stranded mRNA libraries were generated from lung tissues for the male F1 fetus for transcriptomics. Proximo Hi-C libraries were also generated by cross-linking approximately 10 mg of fetal lung tissues using the Proximo Hi-C v4.0 protocol, following the manufacturer's recommendations and sequenced on an Illumina NextSeq2000 sequencer with paired end 2 × 151 cycles. A summary of all the data generated from each technology for all the samples used is provided in Table S1.

Table S1: Summary of the reads generated from the different technologies for the Bighorn male and female F1 samples

| | **Male F1 (coverage)** | **Female F1 (coverage)** |
|---|---|---|
| **HiFi yield** | 172.95 Gb (57x) | 238.6 Gb (83x) |
| **ONT yield** | 714.84 Gb (137x) | 789.28 Gb (141x) |
| **Proximo HiC** | 69.90 Gb (26x) | |

### 3.0 Estimation of Heterozygosity

To increase the success of phasing the diploid genome of the F1 cross a high level of heterozygosity was ensured between the parents. The estimation of the heterozygosity was done by using Jellyfish to count the kmers in the F1 HiFi reads data before running Genomescope[1] v2.0 (Figure S1).

A canonical 21-mer database was produced with Jellyfish:

*jellyfish count* ***-m $kmer -s*** *1000M* ***-t $threads -C -o $outFileName $HiFiReads***

The kmer histogram used as Genomescope input was produced with:

*jellyfish histo* ***-t $threads $outFileName*** > ***$histoOutFile***

Genomescope was run with the following parameters to estimate the heterozygosity

*Rscript genomescope.R* ***$histoOutFile*** *"21" "16000" outGFolder*”

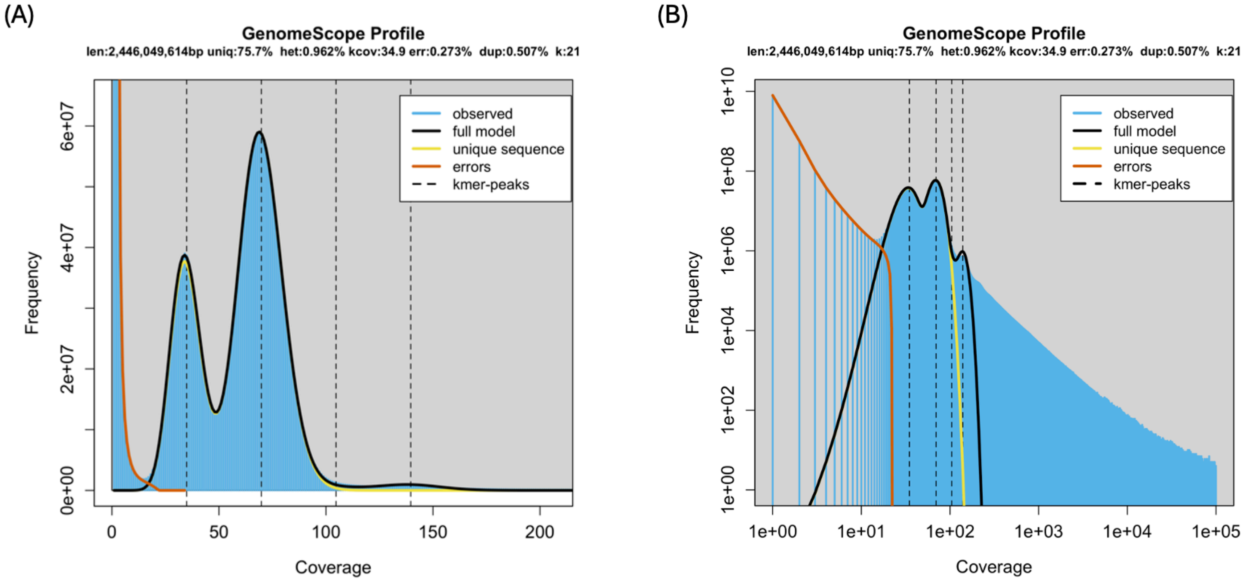

Figure S1: Estimation of heterozygosity in the male F1 cross. (a) The Genomescope model of the raw Illumina reads highlighting the kmer distribution of the reads in the heterozygous and homozygous coverage regions. The estimated heterozygosity is 0.96%. (b) zoomed in view of the model.

### 4.0 Bighorn genome assembly

#### 4.1 Male F1: All autosomes and chromosome Y

Three different versions of the Bighorn draft assembly was produced using the same HiFi, ONT and parental Illumina reads, but with three different versions of Verkko[3] assembler– Verkko_v1.1 (assembly *Bighorn_ASM_v1.1*), Verkko_v1.4 (assembly *Bighorn_ASM_v1.4*) and Verkko_v2.0 (assembly *Bighorn_ASM_v2.0* used for the Bighorn-T2T assembly). *Bighorn_ASM_v2.0* was a 2.83 Gb draft assembly of the paternal bighorn haplotype in 159 contigs with contig N50 of 103.34 Mb (Table S2). This draft assembly comprised 19 single contig T2T chromosomes with telomere sequence at their ends out of the total 27 (26 autosomes and the Y chromosome) (Supplementary Data 1). The remaining chromosomes were subjected to additional manual curation steps described below to bring them to telomere-to-telomere status.

##### 4.1.1 Manual assembly curation

There were two main categories of contig breaks in the draft *Bighorn_ASM_v2.0* assembly that required manual curation to attain T2T status (Figure S2) – alternative ambiguous paths in the assembly graph (also referred to as tangles), and unanchored telomere-bearing contigs.

(i.) The assembly graph tangles resolution, affecting OCA1, OCA4, OCA10 and OCA24, entailed identifying the correct path which is a simple walk through the adjacent nodes in a tangle using a combination of visual inspection of the assembly graph

Table S2: Assembly statistics of the Bighorn male and female Ff1 fetuses

| | Male F1 | | Female F1 | |
|---|---|---|---|---|
| | **Hap1-Polypay** | **Hap2-Bighorn-Male** | **Hap1-Polypay-Female-F1** | **Hap2-Bighorn-Female-F1** |
| No_scaffolds | 947 | 159 | 1,766 | 1,886 |
| Total_length | 2,950,008,371 | 2,836,304,228 | 3,014,546,206 | 3,127,591,003 |
| Average_length | 3,115,109.1 | 17,838,391.3 | 1,706,991.0 | 1,658,319.7 |
| Scaffold_N50 | 96,765,252 | 103,349,577 | 95,201,683 | 103,557,659 |
| Scaffold_auN | 106,999,036.8 | 132,260,062.1 | 123,592,340.2 | 129,429,639.0 |
| Scaffold_L50 | 12 | 9 | 10 | 10 |
| Largest_scaffold | 284,745,837 | 280,946,186 | 283,879,984 | 281,398,027 |
| Smallest_scaffold | 5,228 | 9,062 | 4,103 | 8,013 |
| No_contigs | 954 | 166 | 1,773 | 1,892 |
| Total_length | 2,948,933,313 | 2,835,065,322 | 3,013,105,771 | 3,127,538,420 |
| Average_length | 3,091,125.0 | 17,078,706.7 | 1,699,439.2 | 1,653,032.9 |
| Contig_N50 | 96,765,252 | 103,349,577 | 95,201,683 | 103,557,659 |
| Contig_aUN | 106,421,894.4 | 131,848,086.5 | 121,673,908.1 | 126,395,388.4 |
| Contig_L50 | 12 | 9 | 10 | 10 |
| Largest_contig | 284,745,837 | 280,946,186 | 283,879,984 | 281,398,027 |
| Smallest_contig | 5,228 | 9,062 | 4,103 | 8,013 |
| No_gaps_in_scaffold | 7 | 7 | 7 | 6 |
| Total_gaps_in_scaffold | 1,075,058 | 1,238,906 | 1,440,435 | 52,583 |
| Average_gaps_in_scaffold | 153,579.7 | 176,986.5 | 205,776.4 | 8,763.8 |
| Gap_N50 | 542,528 | 361,747 | 475,967 | 31,583 |
| Gap_auN | 448,415.9 | 392,740.7 | 469,783.2 | 20,890.5 |
| Gap_L50 | 1 | 2 | 2 | 1 |
| Largest_gap_in_scaffold | 542,528 | 527,121 | 485,980 | 31,583 |
| Smallest_gap_in_scaffold | 1,000 | 3,663 | 1,000 | 1,000 |
| GC_content_% | 43.9 | 44.1 | 44.3 | 44.9 |

(i.) with Bandage[4], coverage evidence produced by Verkko, and reads alignments to the assembly graph.

(ii.) Eight chromosomes were assembled in contigs having telomere repeat sequence at only one end, except OCA25 which lacked telomere sequence at both ends (Supplementary Data 1). Contigs of lengths ranging between 66.7kb and 18.08Mb harboring telomere repeat sequence at one of the ends (Table S2) but not anchored on any chromosome were identified in the draft assembly. These contigs (previously referred to as unanchored floating telomeric contigs) clearly belonged with the incomplete chromosomes that were assembled without telomeres at the ends but only required identification of the correct contig (OCA3, OCA5, OCA6, OCA11, OCA12, OCA14, OCA20 and OCA25). Identification and pairing of the floating telomeric contigs with the correct chromosomes were done through various manual curation steps as described below.

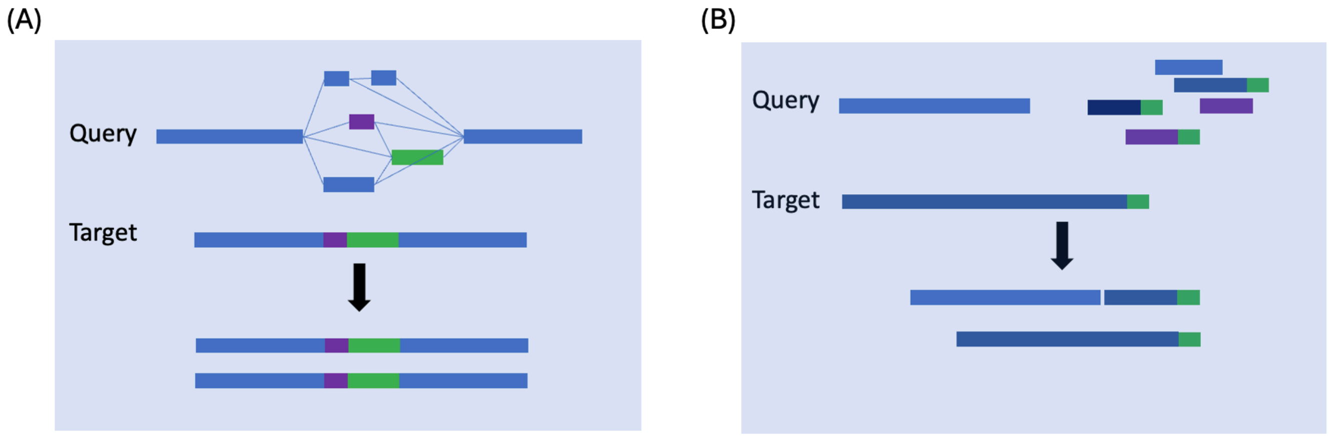

Figure S2: Categories of contig breaks on the Bighorn draft assemblies. (A) Graph tangles being resolved with longer sequence (B) Floating telomeric sequence being anchored on the respective chromosome. The query sequence is the chromosome with either tangles or without one or both telomere ends while the target sequence is the contig being used to guide the alignment. In figure B, the green shade on the contigs are the telomeres.

Table S3: Floating telomere-bearing contigs in the Bighorn male F1 draft assembly

| Contig_ID | telo-start (bp) | telo-end (bp) | Length (bp) |
|---|---|---|---|
| pat-0001239 | 11,094,443 | 11,111,091 | 11,111,091 |
| pat-0001240 | 0 | 22,622 | 12,371,894 |
| pat-0001245 | 12,579,510 | 12,598,783 | 12,598,783 |
| pat-0001247 | 18,062,829 | 18,080,311 | 18,080,311 |
| pat-0001250 | 0 | 17,675 | 14,336,904 |
| pat-0001263 | 1,138,198 | 1,158,408 | 1,158,408 |
| pat-0001269 | 13,508,259 | 13,526,259 | 13,526,259 |
| pat-0001289 | 111,730 | 128,477 | 128,477 |
| pat-0001290 | 70,774 | 87,258 | 87,258 |
| pat-0001314 | 41,425 | 66,742 | 66,742 |
| pat-0001354 | 0 | 20,781 | 128,629 |
| pat-0001358 | 100,665 | 125,864 | 125,864 |
| pat-0001387 | 122,041 | 140,311 | 140,311 |

GraphAligner[5] and Winnowmap2[6] were used to align the ONT ultra-long (UL) reads to the assembly graph. Winnowmap2[6] was used since it has been reported to exhibit superior performance in the repetitive regions over other long reads aligners[7]. The resulting *.gaf* alignment file was used to define the correct path taken by nodes within tangles which are spanned by the UL reads. The paths with reported contiguous nodes were collected and supplied to Verkko to produce new consensus sequence for the chromosomes from the newly defined paths.

In the different versions of the assemblies generated (*Bighorn_ASM_v1.1*, *Bighorn_ASM_v1.4*, *Bighorn_ASM_v2.0*), some chromosomes were more contiguous in a version compared to the others (Supplementary Data 19). These differences were likely due to the differences in the heuristics used by Verkko for tangles resolution in the challenging regions of the genome. For each gapped T2T scaffold in the *Bighorn_ASM_v2.0* assembly, the corresponding region in chromosomes of the *Bighorn_ASM_v1.1* and *Bighorn_ASM_v1.4* assemblies were aligned to the assembly graph using GraphAligner[5] to scaffold the gaps.

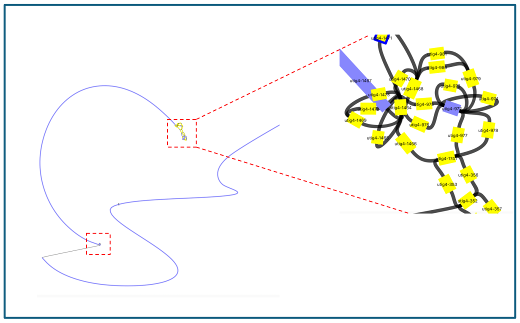

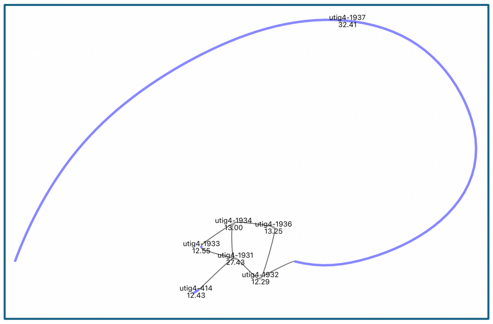

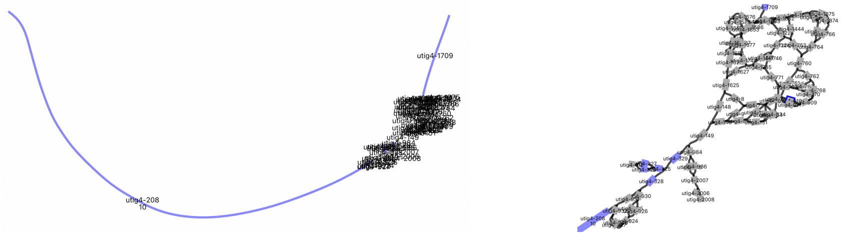

Figure S3: Tangles in the assembly graph of the draft Male F1 fetus assembly on OCA1, OCA6 and OCA10, visualized with Bandage. These tangles resulted in the gaps on the chromosomes due to ambiguous paths for nodes at these loci on the graph.

By using this alignment approach, the tangles on OCA1 (Figure S3), OCA4 and OCA24 were resolved to produce single contiguous T2T chromosomes. A 479.64 Mb gap on OCA10 (Figure S3) was however not amenable to resolution in this manner since no suitable path was identified in the assembly graph to fill the gap. A (peri)centromeric repeat sequence was identified to be enriched at this locus (Supplementary Data 5). The *Bighorn_ASM_v1.1* OCA10 which spanned this gap on *Bighorn_ASM_v2.0* OCA10 with sufficient alignments anchored on the flanks of the gap was used to patch this gap. A new consensus sequence was subsequently produced by Verkko[3] with this sequence.

The missing telomeres on OCA3, OCA5, OCA11 and OCA12 were also successfully identified and joined to the respective floating telomeric contig by aligning the chromosomes to the corresponding *Bighorn_ASM_v1.1* chromosome. The floating telomeric contigs (Table S3) belonging to OCA6 (Figure S3), OCA14, OCA20 and OCA25 however required a different approach to assign them since there were no identifiable paths in the assembly graph to connect them to the respective chromosome.

To resolve these chromosomes, the *Bighorn_ASM_v2.0* assembly was scaffolded with the short reads Hi-C data as extra evidence using the Hi-C mode of Verkko v2.0. Verkko provides estimates of the number of times that all the contigs in the assembly interact with each other from the Hi-C reads alignments, where a higher number is indicative of frequent contact between contigs. This information was used as evidence to anchor the correct floating telomeric contig on OCA6, OCA14, OCA20 and OCA25 to generate new paths. The new paths were subsequently supplied to Verkko to produce new consensus sequence from the nodes defined in the paths.

Post curation checks through alignment of the ONT reads to the assembly revealed some insertions and soft clipping of reads at the junction where the contigs and floating telomere contigs for OCA6, OCA14 and OCA20 were joined (Figure S4). These loci were identified to be (peri) centromeric loci and enriched for repetitive DNA on these chromosomes. The junctions were scaffolded with 100 N’s per NCBI standard to signify unknown copies of repetitive sequence in a genomic region since it was difficult to ascertain the full span of the centromeric repeat breaking the alignment at the loci.

(A)

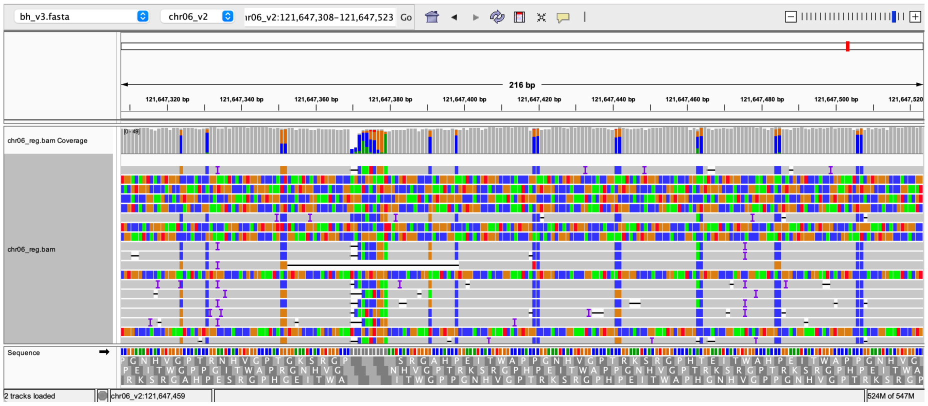

(B)

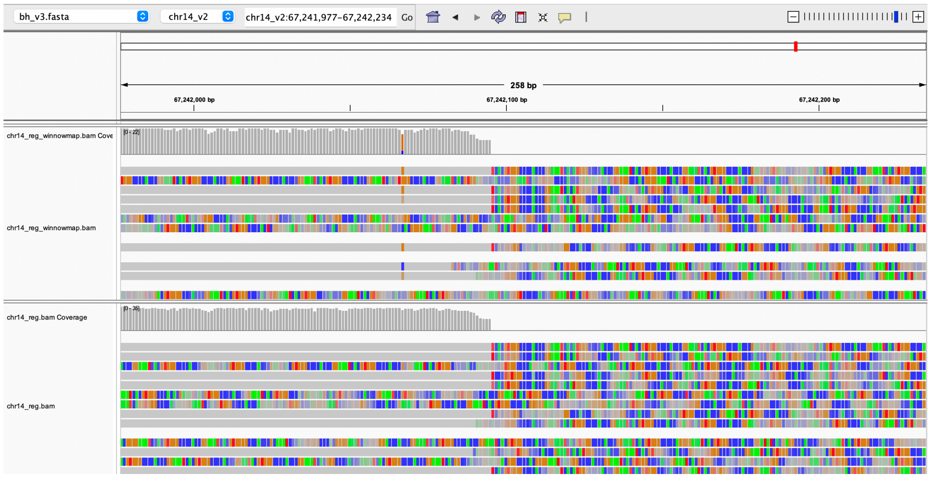

(C)

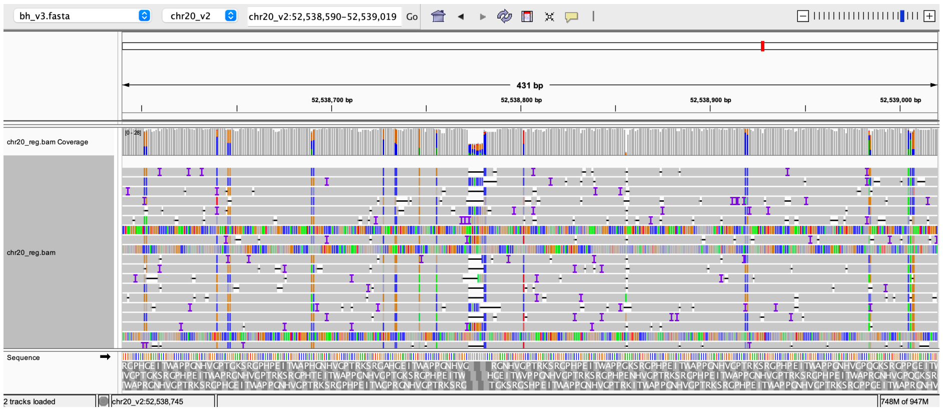

Figure S4: Post curation reads alignment of the ONT reads to the *Bighorn_ASM_v2.0* assembly. IGV view of OCA6 (A), OCA14(B) and OCA20 (C) showing the insertions and soft-clipping at the junction of the chromosomes and the floating telomeric contigs joined to them.

Manual curation of the draft *Bighorn_ASM_v2.0* assembly fixed a total of six contig breaks (two each on OCA4 and OCA24, and one each on OCA1 and OCA10) and attached telomeric contigs to eight incomplete chromosome contigs to make the assembly telomere-to-telomere.

##### 4.1.2 Assembly Polishing

The manually curated Bighorn T2T assembly was validated by checking the concordance between the raw reads used for the assembly and the final curated assembly by mapping all the HiFi and the ONT reads separately to the curated assembly with Winnowmap2[6,8].

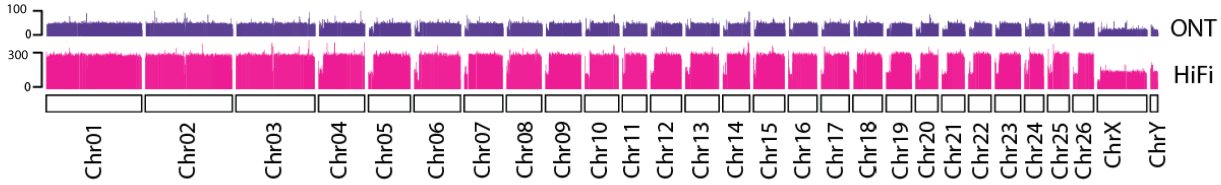

Figure S5: Concordance of the HiFi and ONT reads with the manually curated Bighorn-T2T assembly.

The distribution of the microsatellites AT, GA, GC, and TC across the genome was estimated using the T2T-Polish[9] pipeline to identify potential assembly issues due to known sequencing technology bias[10,11]. T2T-Polish[9] was also used to estimate the coverage of the HiFi and ONT reads (Figure S5) along the entire genome assembly.

The curated bighorn assembly was polished using the F1 HiFi and Illumina short reads independently with the Racon-based T2T-polish pipeline[9] to improve the base accuracy. The T2T-polish pipeline used Winnowmap2[6,8] to align the F1 reads to the assembly, filtered the alignments with Falconc (https://github.com/bio-nim/pb-falconc/releases) before calling variants with Racon (https://github.com/isovic/racon/tree/liftover). The variants were filtered with Merfin[12] while a new consensus genome assembly was produced with Bcftools[13] based on the Merfin[12]-filtered variants call. A hybrid k-mer database of the

Illumina and HiFi reads was used for polishing and base accuracy evaluation to eliminate the effects of sequencing technology biases on k-mers[9].

The high-quality hybrid k-mer database was produced using the following command:

*meryl union-sum [ greater-than 1 F1_31_clean.meryl ] [ greater-than 1 hifi_31.meryl ] output hybrid_31.meryl*

The base accuracy QV value of Bighorn-T2T was improved from 61.77 to 63.22 after four rounds of polishing with the HiFi reads. The Illumina data was then applied to the polishing for marginal increase of the QV to 63.76 (Supplementary Data 3). Alignment of the long reads was done with the repeat-aware aligner Winnowmap2[14] including the -I8g parameter to create large index for genome assemblies that are larger than 4Gbp.

#### 4.2 Female F1: Chromosome X

The bighorn assembly generated from the male haploid of the F1 produced only the Y sex chromosome. A second interspecies cross of a female F1 was sequenced for the X chromosome. The sequence data for the female fetus included 238 Gb HiFi (85x), 128 Gb ONT UL (45x), a total of 126M reads from two Hi-C libraries, and *92 Gb (31x)* and *89 Gb (30x)* of whole genome short reads from dam and sire, respectively. The draft assembly of the sire haplotype of the female fetus had a length of 3.12 Gb and contig N50 of 103.55 Mb (Table S1), with the X-chromosome assembled in a single contig harboring telomere sequence at both ends.  Without taking the whole chromosomes in the whole assembly to

T2T, the assembly was polished and chromosome X was taken to supplement the complete bighorn assembly of the male f1.

### 5.0 Validation of nucleolar organizer region (NOR) *rDNA* array lack on OCA2

Annotation of the Bighorn-T2T assembly revealed that rDNA arrays were located on OCA1, OCA3, OCA4 and OCA25 (Table S4). An array is expected to be located on OCA2 as well, based on the reported presence of an array on domestic sheep OAR2 from Silver staining experiment [15].

Table S4: NOR rDNA array copies and their location on the Bighorn-T2T genome

| **Chromosome** | **rDNA_copies** | **start (bp)** | **end (bp)** |
|---|---|---|---|
| OCA1 | 9 | 78,119 | 385,994 |
| OCA3 | 2 | 232,245,465 | 232,321,593 |
| OCA4 | 12 | 135,229,911 | 135,687,895 |
| OCA25 | 3 | 63,323,479 | 63,435,387 |

This lack of an rDNA array on OCA2 was further investigated using the different approaches described as follows.

The raw ONT reads and the Herro[16]-corrected ONT reads were independently aligned to Bighorn-T2T assembly and the aligned reads of length >= 70kb and bearing telomere sequence at the ends were extracted to be checked for the presence of rDNA sequence. The reads alignment showed sufficient coverage at the end of OCA2 up to the telomeric region (Figure S6). RepeatMasker[17] was run on these reads to identify any rDNA sequence. None of the reads indicated the presence of any rDNA component.

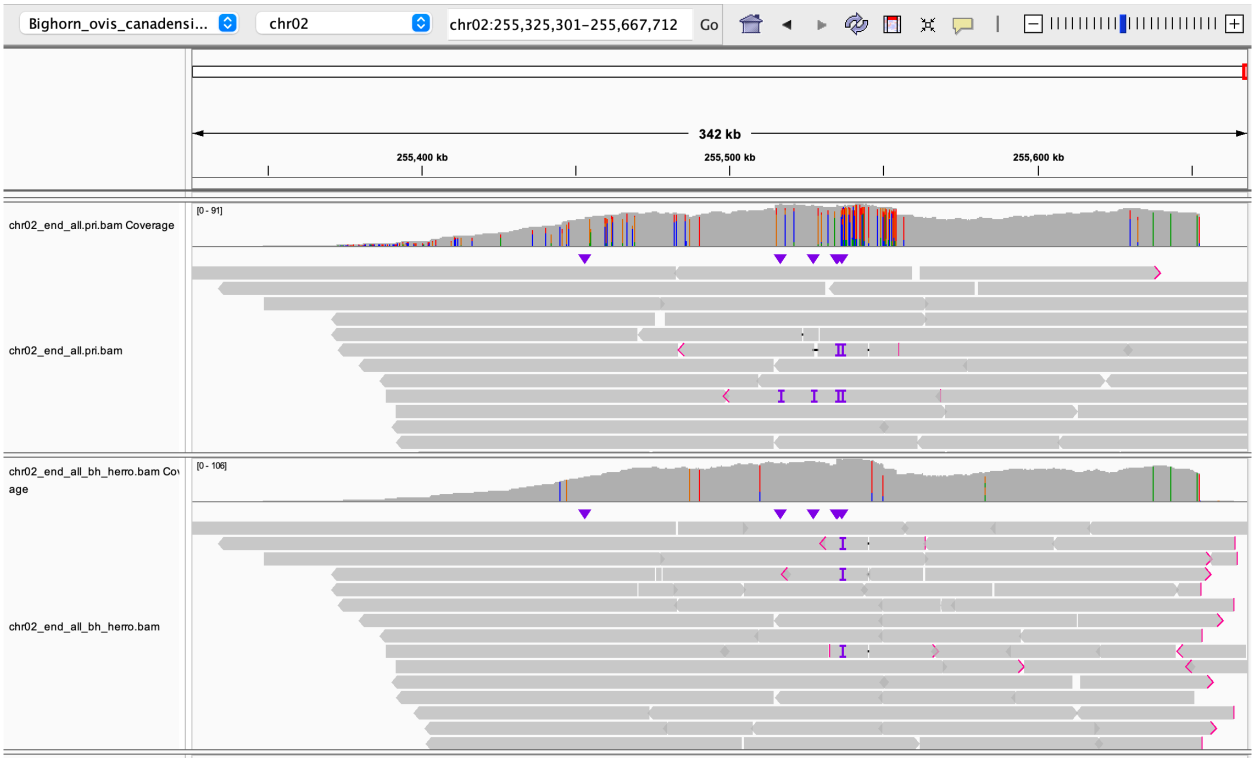

Figure S6: ONT reads alignment to the q-arm end of Bighorn-T2T OCA2. IGV visualization of alignment coverage of telomere-bearing raw ONT UL (top track) and Herro-corrected (bottom track) reads at the q-arm of the Bighorn-T2T OCA2 showing sufficient coverage of the reads terminating the chromosome assembly.

We further investigated the absence of a NOR on OCA2 using other different approaches. OCA2 of the female F1 cross (henceforth FemF1-OCA2) was examined for the NOR locus and was confirmed to contain 2 full copies of rDNA before the contig break. Pairwise alignment was then made with the Bighorn-T2T OCA2 (henceforth Bighorn-T2T-OCA2) to identify the approximate corresponding NOR locus. The last 500kb of the two chromosomes were extracted with *seqtk subseq* for alignment with Minimap2[18]. The alignment exhibited collinearity between the two chromosomes only up to 303.5 kbp on FemF1-OCA2 but up to the start of the telomere sequence on Bighorn-T2T-OCA2 (Figure S7). The rDNA array on FemF1-OCA2 was however located at 102,158 bp into the remaining 196.4 kb that was absent on the Bighorn-T2T-OCA2.

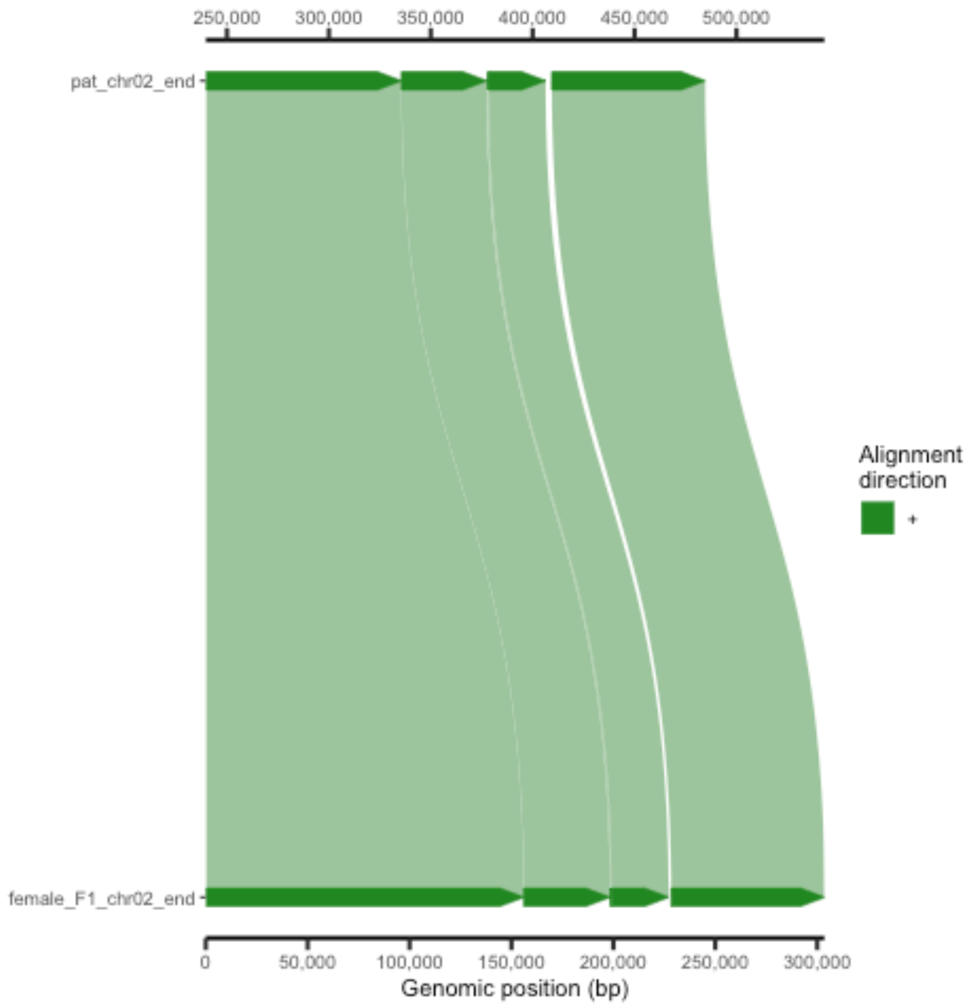

Figure S7: Synteny alignment between the 500kbp ends of Bighorn-T2T-OCA2 and FemF1-OCA2

We further screened the Human 45S rDNA unit (HSA_rDNA henceforth, with accession NR_046235.3 on NCBI) on Bighorn-T2T-OCA2 using Mash screen[19]. Mash screen was run by first creating a sketch of the HSA_rDNA using default parameters before running Mash screen to report all alignments using the option -i -1. The result indicated only 34/1000 were shared between the HSA_rDNA and Bighorn-T2T-OCA2 assembly. The 13.3kb-long HSA_rDNA sequence was aligned to the complete bighorn assembly using Blast. While the longest alignment block on chromosome 2 was just 710 bp (recorded at 82.8Mb) about 168Mb away from the expected NOR region, about 4.1kb alignment block of the HSA_rDNA was consistently recorded for each rDNA unit on OCA1, OCA3, OCA4 and OCA25.

The microsatellites tracks generated for the bighorn assembly after manual curation was also examined around the expected NOR locus on Bighorn-T2T-OCA2 and on the other chromosomes containing the NOR for indications of any truncation of the assembly in this region. The clearly defined undulating patterns in the GC microsatellites content observed at the NOR loci on the other NOR-bearing chromosomes was conspicuously absent on Bighorn-T2T-OCA2 (Figure S8).

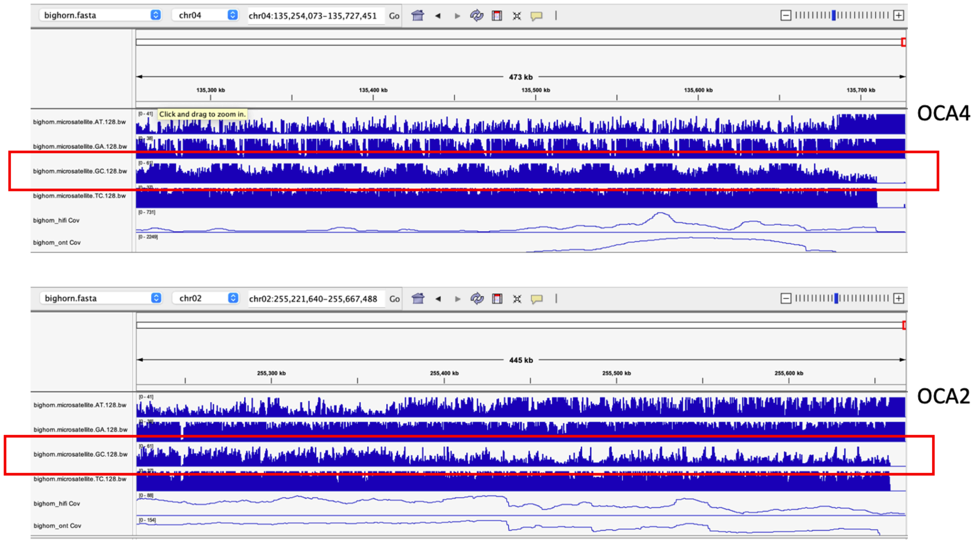

Figure S8: Tracks showing the AT, GA, GC and TC microsatellites distribution within 128bp windows at the NOR of OCA4 and the expected NOR locus on OCA2.

Combining all this evidence led us to conclude that an rDNA array was absent from OCA2 of the male F1 used to produce Bighorn-T2T.

### 6.0 Repetitive sequence annotation

#### 6.1 Ovine and bovine satellites annotation

The DNA sequence of the known ovine satellite 1.714 (Accession: X01839.1) and the bovine satellites[21,22] were obtained from NCBI and aligned to Bighorn-T2T using the Blast[23] program.

*blastn -task blastn -query $1 -db $2 -out $3*

where $1 is the query sequence, $2 is the bighorn blast database and $3 is the output file.

The (peri)centromeric regions of all chromosomes were observed to be enriched for the bovine satellites Sat1.723 and Sat1.706 (co-located but interleaved in opposite orientation) as well as the ovine satellite 1.714 in different combinations, except the Y-chromosome which was enriched with a different repeat sequence we had previously identified on the domestic sheep OARY[24]. The enrichment of different combinations of bovine satellites families observed at the (peri)centromeric region of all the bighorn chromosomes is well documented in bovine genomes[21,22]. The span of centromeric satellites were much shorter on the submetacentric chromosomes OCA1, OCA2 and OCA3 compared to the acrocentric chromosomes. This observation agrees with previous reports of the reorganization of centromeric DNA through loss of or reduction of satellites when metacentric chromosomes were formed by ancient Robertsonian translocations on sheep[25] and other mammals[26,27].

#### 6.2 (Peri)centromeric repeats

While manually curating the assembly to join chromosomes without telomere sequences (described in manual curation section above), OCA6, OCA14 and OCA20 contigs appeared broken at a repetitive sequence. Manual decomposition of the sequence at these loci revealed a 20bp monomeric sequence which was arranged into a 242bp higher-order repeat (HOR) comprising the monomeric sequence separated by dinucleotides (Main Figure 2). To estimate the abundance of this sequence across the genome, the HOR sequence was aligned to Bighorn-T2T with Blast[23] while the result was filtered for alignment coverage of at least 80% between the sequence and the target. This sequence was observed to be enriched at the centromeres of most of the chromosomes (Figure S9). This HOR sequence added 31,179,308 bp of sequence to the Bighorn-T2T genome and accounted for about 4.3% of the NARs.

20bp Monomer sequence: GGAAATCACGTGGGCCCCAC

242bp HOR sequence:

AGGGAAATCACGTGGGCCCCACCCGGAAATCACGTGGGCCCCACCCGGAAATCACGTGGG

CCCCACCAGGAAATCACGTGGGCCCCACCAGGAAATCACGTGGGCCCCACCCGGAAATCAC

GTGGGCCCCACCCGGAAATCACGTGGGCCCCACAGGGAAATCACGTGGGCCCCACCCGGA

AATCACGTGGGCCCCACCCGGAAATCACGTGGGCCCCACCAGGAAATCACGTGGGCCCCAC

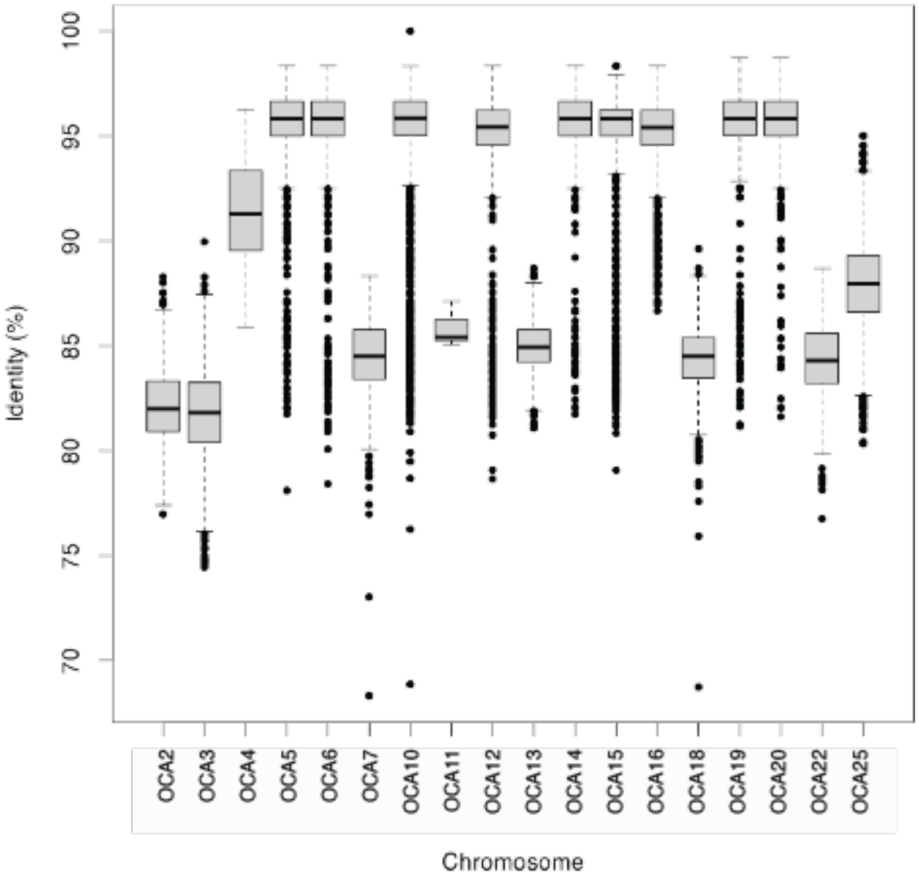

Figure S9: Sequence identity of the copies of the newly identified (peri)centromeric HOR repeat on the bighorn genome assembly. There is remarkable difference in sequence identity between the copies of the repeat unit between the chromosomes; the submetacentric OCA2 and OCA3 generally had lower sequence identity alongside some of the acrocentric chromosomes.

Using the same approach, the abundance of the HOR sequence was estimated on the Bighorn-v1 assembly (GCA_004026945.1 on NCBI), as well as on the domestic sheep reference assembly[28]. Since the bovine satellite classes have been reported to share some degree of sequence similarity between them[22], the HOR was checked on the Ovine Sat 1.714 and the Bovine satellites, but no sequence homology was observed.

#### 6.3 RepeatMasker annotation

RepeatMasker[17] v4.1.4 was run with RMBlastn v2.13.0+ using –species "*sheep*" to obtain a general sense of the distribution of repetitive elements (RE) across Bighorn-T2T. The genome was also hard-masked for the downstream steps in the *de-novo* RE discovery pipeline (Supplementary Data 9). The annotation result was filtered for simple repeats and alternative alignments prior to further analysis with

*grep -v "*" bighorn.fasta.out | grep -v "Simple_repeat" | awk 'OFS="\t"{strand="+"; if($9=="C") strand="-"; {print $5,$6,$7,$10"+"$11,$2,strand}}' | tail -n+4 > RM_less_simple_R.bed*

#### 6.4 De-novo repeats discovery and annotation

RepeatModeler[29] v2.0 was run on the hard-masked genome to facilitate identification of new repeat models in the regions that were not annotated by RepeatMasker[17]. Curation of the repeat models produced by RepeatModeler were carried out using Tetrimmer[30] (https://github.com/qjiangzhao/TEtrimmer) and Earl Grey[31] by automating the process of repeat model extension and consensus building with their pipelines. The two tools use the consensus models from RepeatModeler as input but were run independently.

*Python3 **Tetrimmer.py** –input_file **$inFile** –genome_file **$ASM** –output_dir **$outDir** –pfam_dir **$pfam** –classify_all* where $inFile is the fasta file containing the consensi repeat models from RepeatModeler, $ASM is the genome assembly file, $pfam is the path to the Pfam database and the classify_all parameter tells the pipeline to classify every consensus sequence.

Similarly, EarlGrey was run as follows:

*earlGrey* ***-g $inFile -s $species -o $outDir -r $rmSearch -c*** *yes* ***-m*** *yes* ***-d*** *yes*

where $inFile is the genome assembly, $rmSearch is the repeatmasker search term ("sheep" was used), and the parameters -c to cluster the TE library to reduce sequence redundancy, -m to remove TE annotations less than 100bp and -d to produce a soft-masked genome when the analysis is completed.

One of the final repetitive element annotations produced by TEtrimmer[30] is a 23,204bp annotated as a Long Terminal Repeat/Endogenous Retroviruses (LTR/ERV1) element although without the expected terminal LTR sequence (Figure S10). TEtrimmer extended this element from a 621bp consensus sequence (RepeatModeler repeat models) and it subsumes a tandem array of the novel 242bp novel repeat described in the (Peri)centromeric repeat section above. This 23kb LTR/ERV element is structured into a

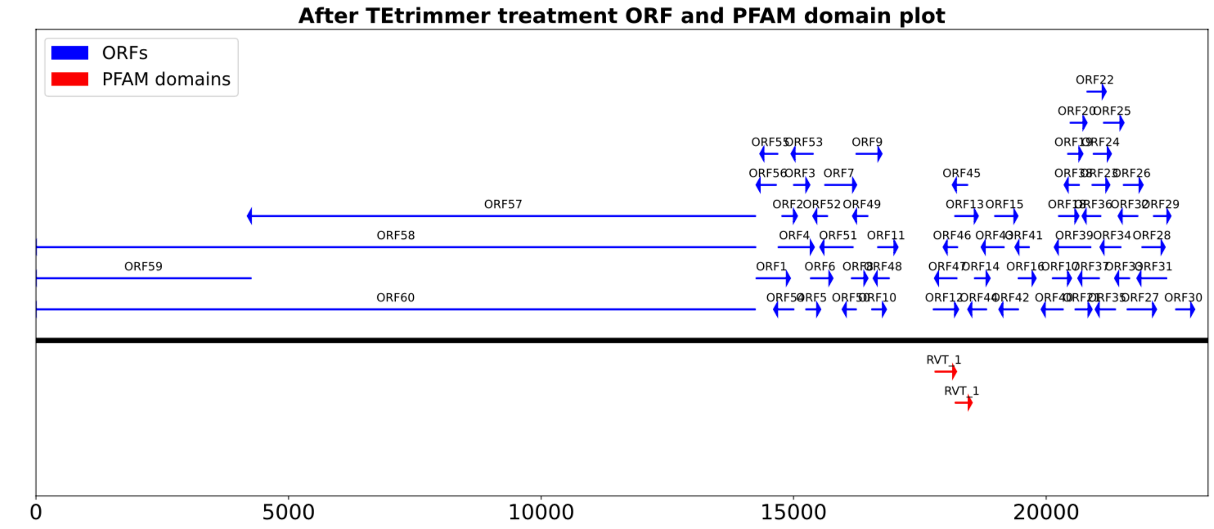

Figure S10: TEtrimmer annotation of the 23kb LTR/ERV element on the Bighorn-T2T genome

tandem repeat of the centromeric HOR spanning about 14.2kb (segment1) and the remaining 8.9kb (segment2) lacks this repeat sequence; both segments were separated by 420bp (Figure S11). This LTR/ERV element was also checked on the Cattle (*Bos taurus*), Sheep (*Ovis aries*), Goat (*Capra hircus*), Argali (*Ovis ammon*), and Mouflon (*Ovis orientalis*) genomes. Although the Argali and the Mouflon genome assemblies that were used to check the presence and abundance of the LTR/ERV element were not T2T, the two segments of the LTR/ERV element were found to be enriched on these genomes (Figure S12). Only Bighorn-T2T recorded full length copies (25) of the 23kb ERV/LTR element (Figure S12). The absence of this repeat model on a cattle T2T genome (GCA_040286185.1 on NCBI) is suggestive of emergence in sheep after the split from cattle about 19MYA[33] (Figure S12). Its reduced abundance and sequence identity on goats, which split from sheep about 5MYA[32], could be an indication that it is being lost.

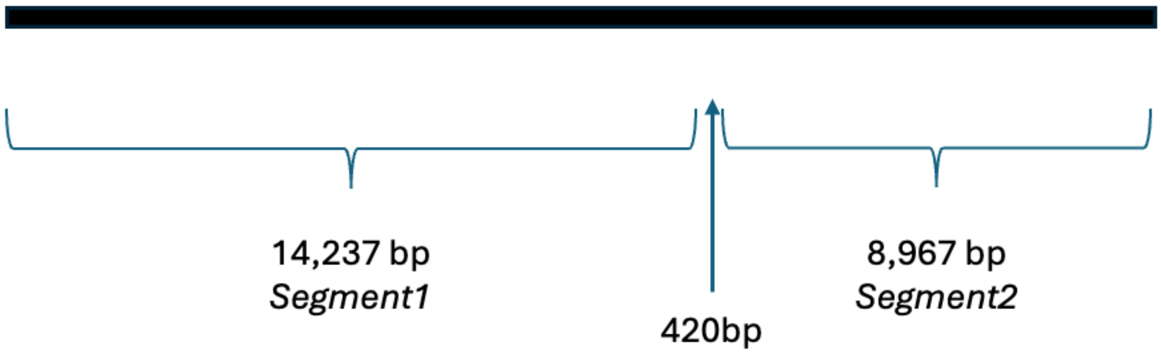

Figure S11: The 23kb repeat element annotated as an LTR/ERV by TEtrimmer comprising 2 segments, segment 1 with the HOR, and segment 2 without it.

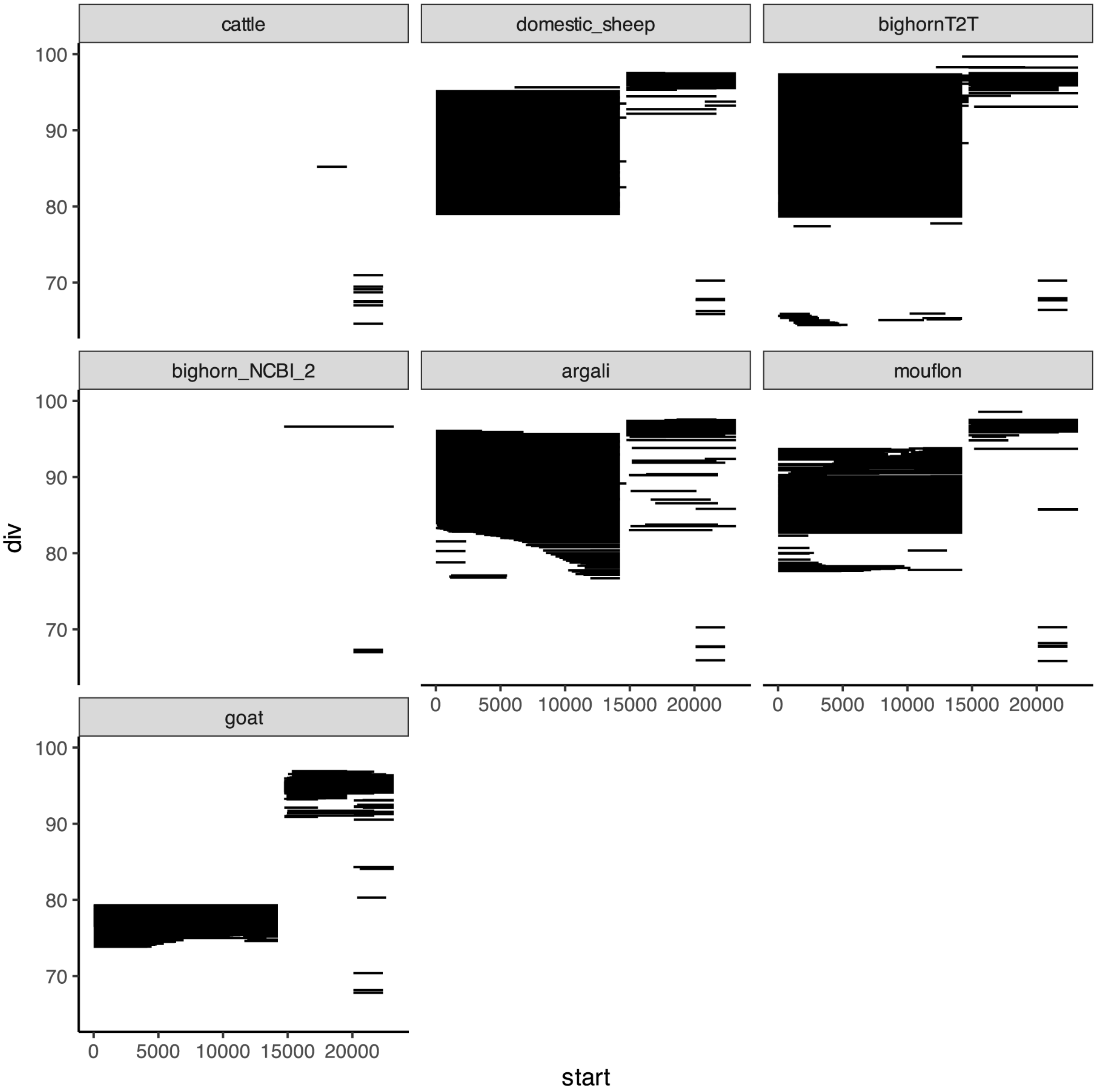

Figure S12: The reads alignment coverage of the 23kb LTR/ERV element showing the span and sequence identity in Cattle, domestic sheep, Bighorn (Bighorn-T2T and Bighorn-v1), Argali, Mouflon and Goat assemblies.

### 7.0 Structural difference analysis

Structural difference (structural variation SVs) between the Bighorn-T2T and the domestic sheep reference assembly[28] were called using PBSV (2.10.0) (https://github.com/PacificBiosciences/pbsv) and Minimap2[18] plus its *Paftools.js* module.

The Bighorn HiFi reads were trio-binned with Canu[37] prior to alignment.

Canu was run as follows:

```
canu -haplotype \
 -p bh_pp_trio \
 -d trioBin \
 genomeSize=3g \
 -haplotypeBighorn illumina_data/sire/*.fastq.gz \
 -haplotypePolypay illumina_data/dam/*.fastq.gz \
 -pacbio-hifi hifi_data/*.fastq.gz
```

#### 7.1 PBSV variants call

Variants calling using pbsv was done by first aligning the Bighorn HiFi reads to the domestic sheep reference assembly using pbmm2 followed by pbsv discover and pbsv call.

```
pbmm2 index $ref "ramb2.mmi"
pbmm2 align --sort --preset  HiFi -j $threads "ramb2.mmi" $inReads
pbsv discover ${inASM}.bam $sample".svsig.gz"
tabix -c '#' -s 3 -b 4 -e 4 $sample".svsig.gz"
pbsv call $ref $sample".svsig.gz" $sample"_pbsv.vcf"
```

where $ref is the domestic sheep reference assembly, $inReads are the Bighorn HiFi reads, $inASM is the bam alignment file from the pbmm2 alignment step and $sample is the output filename of the pbsv discover step.

#### 7.2 Minimap2 variant calls

```
minimap2 -c --cs -L \
 -t $threads ${ref}.mmi ${path}/${query}.fasta \
 | sort -k6,6 -k8,8n > ${tag}_minimap.paf
```

```
k8 paftools.js call -f $ref -s $query -L2000 ${tag}_minimap.paf >
${tag}_minimap_call.vcf
```

#### 7.3 Variants effect prediction with SNPEff

The variants effects predictor SNPEff[38] was used to annotate the variants call (VCF) file to predict the effects of the structural differences on genes. The impact of the variants effects were categorized into HIGH, LOW, MODERATE and MODIFIER based on the changes that these have on the protein-coding gene structure. SNPEff was run as follows:

```
java -Xmx2G -jar snpEff.jar build -gff3 -v chsheept2t
java -Xmx8g -jar snpEff.jar $genome $inVCF >
${tag}_annotated_variants.vcf
```

*where $genome is the ID of the genome in the SNPEff database to be annotated and $inVCF is the input VCF file containing the variants to be annotated*.

### 8.0 Newly assembled genes

To identify the newly assembled genes in bighorn, the gene annotation on the previous assemblies and Bighorn-T2T were compared. Due to a lack of gene annotation, the older bighorn sheep assemblies on NCBI, GCA_004026945.1(Bighorn_v1) and GCA_001039535, were annotated with Bighorn-T2T using Liftoff.

Liftoff was run as follows:

```
liftoff -o $outFile -u $unmapped -flank 0.0 \
    -copies -sc 0.98 -f $features -cds -p $threads \
    -m minimap2 -g $refGFF $targetFile $referenceFile
```

The requirement to enforce matching flanking sequence to the gene was relaxed (-flank 0.0) as well as to allow partial match (by omitting the –exclude-partial flag) to accommodate the limitations of the highly fragmented assembly GCA_004026945.1.

Liftoff annotation showed that 1,991 and 3,701 genes were unmapped to the Bighorn_v1 and GCA_001039535.1, respectively (Table S5). Further manual checks of these umapped genes were carried out by Blast alignment. The tRNAs were excluded from these set of unmapped genes and the rest (499 and 1,589 genes for Bighorn_v1 and GCA_001039535.1) were extracted and aligned to each of Bighorn_v1 and GCA_001039535.1. Blastn was used for the gene alignment to the assemblies with a minimum of 60% coverage between the query and the target.

From the manual gene alignment, a total of 377 (out of 499) and 488 (out of 1,589) genes aligned to Bighorn_v1 and GCA_001039535.1, leaving 122 (Bighorn_v1) and 1,098 (GCA_001039535.1) genes unaligned. The unaligned gene set constitute the genes that were previously not observed on Bighorn_v1 and GCA_001039535.1 assemblies and are newly assembled on Bighorn-T2T.

Table S5: The number of previously unassembled genes on the older bighorn assemblies on NCBI, GCA_004026945.1 (Bighorn_v1) and GCA_001039535.1

| | Liftoff annotation | | | | |
|---|---|---|---|---|---|
| | Number of umnapped genes from Liftoff | | | Manual genes alignment | |
| Assembly accession | All unmapped genes | less tRNA | | aligned | unaligned |
| GCA_004026945.1 | 1,991 | 499 | | 377 | 122 (34 p-coding, 88 non-coding) |
| GCA_001039535.1 | 3,701 | 1,589 | | 488 | 1,098 (538 p-coding, non-coding 560) |

### 9.0 Evaluating the biological significance of a bighorn T2T assembly for population studies

We obtained short-reads genome-wide association studies (GWAS) dataset on NCBI (BioProject ID PRJNA454718, SRA accession SRP144608). This data was from a study [43] which investigated the genetic drivers of *M. ovipneumoniae* carrier status in Bighorn sheep. This dataset was generated using restriction enzyme-based sequencing (Rad-Seq). The original dataset[44] comprised 82 individuals but Martin et al.[43] used a subset of 52 individuals for which longitudinal data on the *M. ovipneumoniae* carrier status data was available. In the original study, the domestic sheep reference assembly Oar_v4.0[45] (GCA_000298735.2 on NCBI) was used. The steps in the analysis pipeline used in the original study were replicated to evaluate the concordance of our results with theirs. Some slight differences were however encountered in the dataset for this analysis compared to the original study – the identifiers of the 52 samples with carrier status data in the original study were not provided. In addition, 18 samples obtained from the same individuals within the population were merged and the identifiers of these samples were not indicated as well. The only phenotypic data (pathogen carriage status) provided was for the final 25 individuals that survived the different data filtering criteria applied in the study.

We filtered our starting dataset from the initial 82 using only the metadata reporting the sex of the individuals in the samples as reported in the paper[43]. The initial dataset was reduced to 61 and this was our input for the analysis. The same analysis steps from the original study were carried out on the same subset of 61 individuals from the original dataset[46] but using Bighorn-T2T as the reference. The population structure was checked (Figure S13) for any stratification that needs to be accounted for in the analysis model. The analysis steps reported[43] was started with Rad-Seq reads processing with STACKS v2.2, end-to-end reads alignment with Bowtie2 (including -sensitive -X 900 parameters) against the domestic sheep *Oar_v4* reference assembly, alignment filtering with SAMtools v1.9[13] using mapping quality > 40 followed by variant discovery and genotyping by STACKS v2.4. The next SNP loci filtering step by minimum depth >=5 and minimum quality score >=20 was also not clear since quality score (QUAL) value is not one of the reported parameters in the output of the STACKS populations program.  The GQ value was used as the parameter reported for this filtering step.  Filtering with PLINK v2 based on the level of missing data per site, minor allele frequency and missing genotypes per sample, linkage disequilibrium (LD) filtering and deviation from the Hardy-Weinberg equilibrium (HWE) were carried out the same way they were reported. The variants from the 56 samples which survived all the filtering criteria (Table S4) were finally subset to keep only the same final 25 individuals with pathogen carrier status phenotypic data used in the original study. A total of 43,020 SNPs remained for association studies after all filtering steps using the Bighorn-T2T as reference genome (Table S4).

(A)

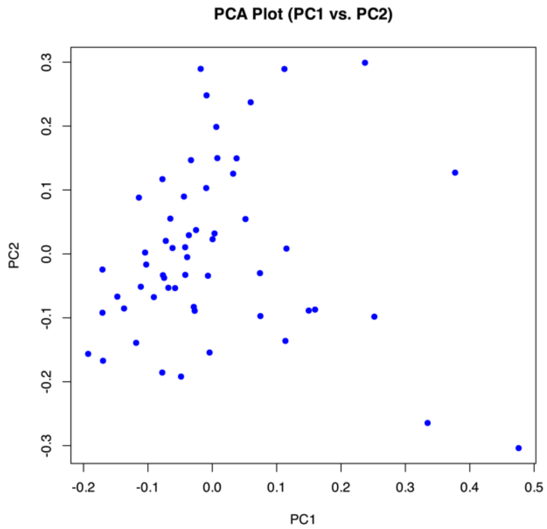

(B)

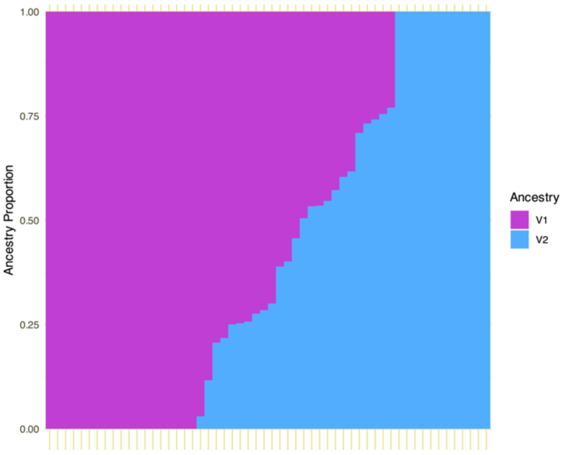

Figure S13: The population structure of the 61 animals used in this study. (A) PCA plot to assess the population structure. (B) Admixture of the population showing the ancestry proportion. Since the individuals are from the same population, the plot has been sorted based on the proportion of ancestry in each individual. Each bar on the x-axis represents an animal.

Due to the stringency of Bonferroni method of p-values correction, we applied a false discovery rate of 0.15 (same as was applied in the original study) using the Benjamini-Hochberg method to control for type I errors.

Table S4: Comparison of the results for the filtering steps in Martin et al.[43] and our reanalysis.

| | Martin et al. (2021) | | | This study | |
|---|---|---|---|---|---|
| | N individuals | N loci | | N individuals | N loci |
| Align individuals to reference genome | 52 | - | | 61 | |
| Minimum quality score>=20 and minimum depth>=5 | 52 | 98,307 | | 61 | 632,928 |
| Remove sites with >10% missing data and MAF 1% | 52 | 17,682 | | 61 | 102,420 |
| Remove samples with >10% missing data | 41 | 17,682 | | 56 | 102,420 |
| LD filtering | 41 | 11,890 | | 56 | 61,976 |
| HWE filtering | 41 | 11,890 | | 25 | 52,050 |
| Remove samples based on bimodal age distribution | 25 | 11,890 | | - | 52,050 |
| GEMMA default filtering | 25 | 10,605 | | 25 | 43,020 |

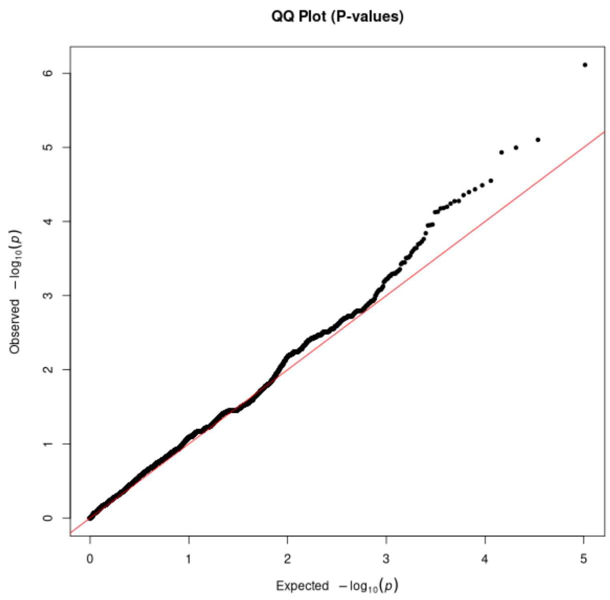

Figure S14: Q-q plot of the SNPs from the reanalysis of bighorn *M. ovipmneumoniae* carrier status study using the bighorn T2T as reference. (The Manhattan plot is on Figure 7 of main section).

### 10.0 Self-identity dotplots

Moddotplot[47] was used to generate the self-dotplots of the chromosomes at 85% sequence identity threshold using the following code;

*moddotplot -i ${inputfile} --no-bed --identity 85 -r 800*
